# Specific vulnerability of long telomeres to undergo end fusions revealed by mutational analysis of Rap1

**DOI:** 10.1101/2021.12.16.472883

**Authors:** Lili Pan, Duncan Tormey, Nadine Bobon, Peter Baumann

**Author notes:** Correspondence, Mailing Address: Department of Molecular Biology, Hanns-Dieter-Hüsch-Weg 15, D-55128 Mainz, Germany. 19396 Piute Cir Bend, OR 97702.

## Abstract

The conserved Rap1 protein is part of the shelterin complex that plays critical roles in chromosome end protection and telomere length homeostasis. Previous studies addressed how fission yeast Rap1 contributes to telomere length maintenance, but the mechanism by which the protein inhibits end fusions has remained elusive. Here, we use a genetic screen in combination with high throughput sequencing to identify several amino acid positions in Rap1 that have a key role in end protection. Interestingly, mutations at these sites render cells susceptible to genome instability in a conditional manner with longer telomeres being prone to undergoing end fusions, while short telomeres are sufficiently protected. The protection of long telomeres requires their nuclear envelope attachment mediated by the Rap1-Bqt4 interaction. Our data demonstrates that longer telomeres pose an additional challenge for the maintenance of genome integrity and provides an explanation for a species-specific upper limit in telomere length.

## Introduction

Telomeres, the structures found at the ends of linear chromosomes, are critical for the maintenance of genome integrity. The leading strand of telomeric DNA is comprised of short G-rich repeats synthesized by the specialized reverse transcriptase telomerase (Pfeiffer & Lingner, 2013). While telomerase solves the end replication problem, telomeres also must be distinguished from DNA double strand breaks. Defects in chromosome end protection results in the activation of a DNA damage response (DDR), which triggers illegitimate repair events including resection and chromosome end-to-end fusions (Palm & de Lange, 2008). Fused chromosomes lead to genomic instability through breakage-fusion-bridge cycles as well as more immediate, massive rearrangements called chromothripsis (Maciejowski *et al*, 2015). To avoid such catastrophic events, telomeric DNA is capped by several proteins collectively referred to as the shelterin complex, which protects telomeres from unwarranted repair events (Palm & de Lange, 2008).

In the fission yeast *Schizosaccharomyces pombe*, the shelterin complex is comprised of five proteins: Taz1 binds double-stranded telomeric repeats and recruits Rap1 (Chikashige & Hiraoka, 2001; Cooper *et al*, 1997; Kanoh & Ishikawa, 2001); whereas Pot1, in complex with Tpz1, binds the single-stranded 3’ overhangs (Baumann & Cech, 2001; Miyoshi *et al*, 2008). These two subcomplexes are bridged by Poz1 (Miyoshi *et al*., 2008). An orthologous molecular bridge, connecting the double-stranded and single-stranded parts of telomeres, is formed by the TIN2 protein in mammalian cells, where the subcomplexes on double-stranded DNA (TRF1, TRF2, RAP1) and single-stranded overhangs (POT1 and TPP1) also share fundamental similarities with their fission yeast counterparts (Baumann & Cech, 2001; Bilaud *et al*, 1997; Broccoli *et al*, 1997; Chong *et al*, 1995; Houghtaling *et al*, 2004; Kim *et al*, 1999; Li *et al*, 2000; Liu *et al*, 2004; Ye *et al*, 2004; Zhong *et al*, 1992). Deletion of *taz1, rap1* or *poz1* in *S. pombe* results in massive telomere elongation, suggesting a role of these factors in inhibiting telomerase (Chikashige & Hiraoka, 2001; Cooper *et al*., 1997; Kanoh & Ishikawa, 2001; Miyoshi *et al*., 2008). Deletion of *taz1* and *rap1* also causes chromosome end-to-end fusions when cells are arrested in G1, supporting a direct role for these factors in end protection (Ferreira & Cooper, 2001; Miller *et al*, 2005). Our previous analysis of Rap1 and Poz1 revealed that their function in telomere length regulation is limited to forming part of a molecular bridge connecting the double- and single-stranded parts of the telomeres and that the entire proteins can be replaced by a short covalent linker between Tpz1 and Taz1 (Pan *et al*, 2015). While such a ‘mini-shelterin’ restores wildtype telomere length, chromosome ends are still undergoing fusions via illegitimate repair events when Rap1, but not Poz1, is absent. Adding back Rap1 restored full protection, indicating that Rap1 prevents end fusions independent of its role as molecular bridge connecting Taz1 and Poz1. A function for Rap1 in inhibiting fusions is also seen in budding yeast and human cells (Bae & Baumann, 2007; Pardo & Marcand, 2005; Sarthy *et al*, 2009), although in humans, this function is redundant with other protective mechanisms and is only observed when TRF2 is absent or telomeres are critically short (Lototska *et al*, 2020; Sarthy *et al*., 2009; Sfeir *et al*, 2010).

The N-terminal half of fission yeast Rap1 contains three domains, a BRCA1 C-terminal (BRCT), a Myb- and a Myb-Like domain (Chikashige & Hiraoka, 2001; Kanoh & Ishikawa, 2001). The C-terminal half harbors a Poz1 interaction domain (PI, (Pan *et al*., 2015)) and a structurally conserved Rap1 C-terminus (RCT), which interacts with Taz1 in *S. pombe*, TRF2 in human, and Sir3 in *S. cerevisiae*, respectively (Chen *et al*, 2011; Li *et al*., 2000; Moretti *et al*, 1994). The PI and RCT domains are separated by a mostly unstructured region of approximately 150 amino acids. In this study, we combined a deletion scan and random mutagenesis screen to identify elements in Rap1 that are critical for preventing telomere fusions. Analysis of the mutagenesis screen by next-generation sequencing identified several amino acid positions in the region between PI and RCT domains that are important for end protection. Intriguingly, mutations at these positions specifically rendered elongated telomeres prone to end fusions, whereas telomeres of wildtype length remained protected in the mutants. Contrary to the conventional thinking that critically short telomeres face the greatest threat of being mistaken for DNA breaks, our data suggests that under certain conditions long telomeres are more likely to undergo end fusions and thus cause genomic instability. These observations provide a conceptional framework for why cells maintain telomeres within a clearly defined size range and why long telomeres may undergo rapid deletion events (Li & Lustig, 1996).

## Results

### The region between PI and RCT domains is important for inhibiting end fusions

To test which part of Rap1 is required for end protection, we created a series of Rap1 domain deletions (Figure 1A). Constructs lacking the BRCT, Myb or Myb-L domain were found to retain the ability of the wildtype protein to prevent end fusions (Figure 1B, lanes 2-4). The ΔPI mutant was previously shown to have elongated telomeres due to the disruption of the interaction between Rap1 and Poz1 (Pan *et al*., 2015). Despite the long telomere phenotype, the ΔPI mutant was proficient at preventing end fusions (lane 5), consistent with previous results that Poz1 is not required for end protection (Pan *et al*., 2015). Among the deletion mutants tested here, only Rap1ΔRCT displayed a chromosome end fusion phenotype (lane 6), consistent with the role of the RCT domain in mediating the interaction with Taz1 and thus being responsible for the recruitment of Rap1 to telomeres (Chen *et al*., 2011). The less severe fusion phenotype observed in the ΔRCT mutant compared to *rap1Δ* (compare lanes 6 and 7) may be due to residual recruitment of Rap1ΔRCT to telomeres through the interaction with Poz1. To further characterize the requirements for end protection, we deleted fragments of increasing size from the N-terminus of Rap1. A fragment comprising amino acids 440 to 693 was sufficient for preventing end fusions (Figure 1C, lane 4), confirming previously reported results (Fujita *et al*, 2012). In contrast, expression of amino acids 513 to 693 or a shorter fragment failed to prevent end fusions similar to *rap1Δ (lanes 6 and 7)*. Amino acids 491-693 displayed a weaker fusion phenotype. This was surprising as the fragment only lacks the PI domain relative to 440-693 and *rap1*ΔPI did not show a fusion phenotype (compare lanes 5 in Figure1B and 1C). One possible explanation for this observation could be that the truncation of amino acids 1-490 affects the folding of the remaining protein sequence. In addition, we cannot exclude the possibility that differences in the relative expression levels is a contributing factor. Western analysis using antibodies against the C-terminal V5-epitope tag present on all constructs showed substantially lower protein levels for Rap1_491-693 compared to full length Rap1 or Rap1_440-693 (Figure S1). It must be noted however, that Rap1_440-693 level is also lower than full length Rap1, yet this deletion mutant is fully functional in the same background and under the same growth conditions.

**Figure 1.**
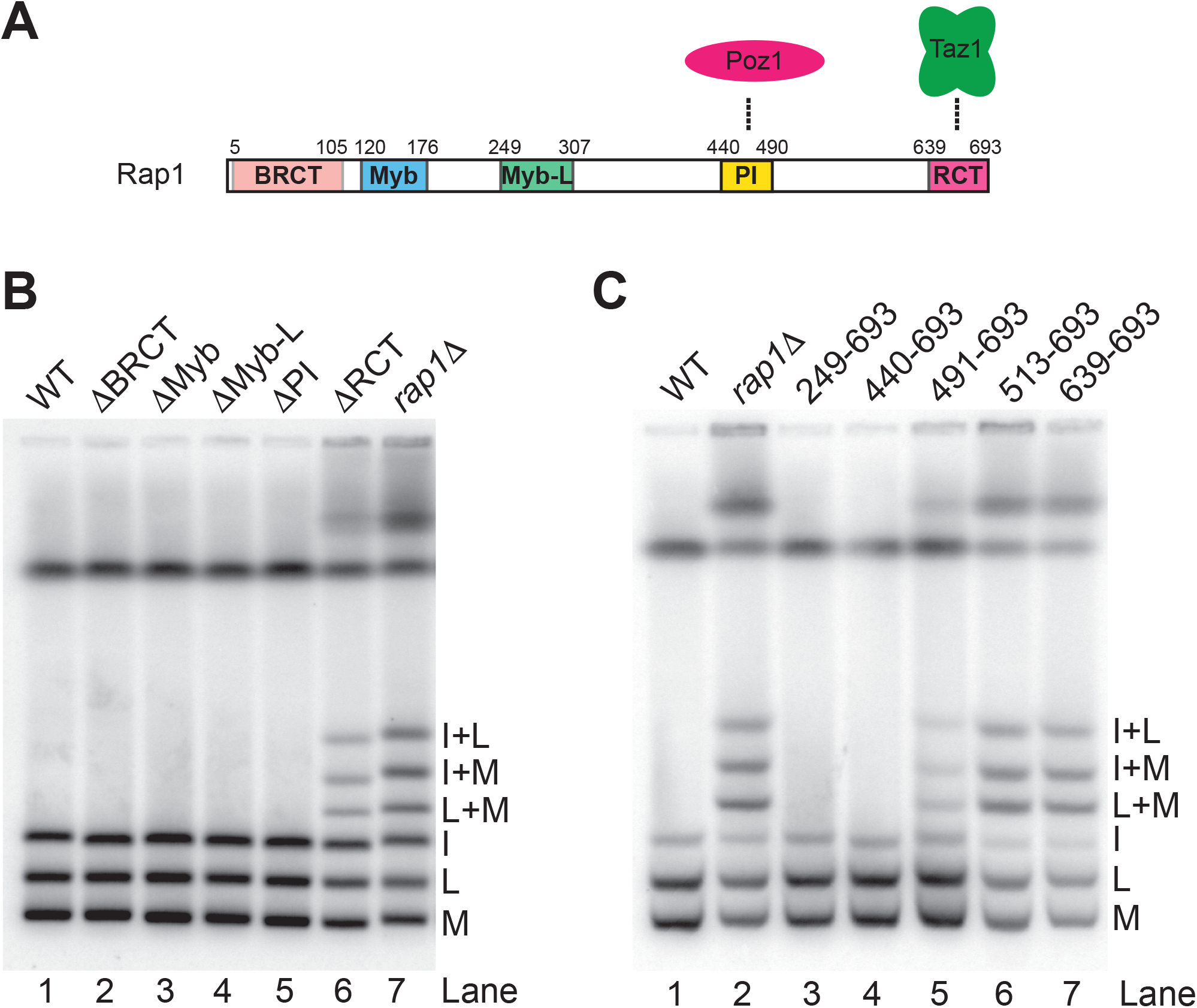
The C-terminal amino acids 440-693 of Rap1 are sufficient to protect telomeres against end fusions. (A) Schematic representation of Rap1 domain structure. (B) End fusion analysis for domain deletion mutants of Rap1. NotI-digested genomic DNA from nitrogen-starved cells was analyzed by pulsed-field gel electrophoresis (PFGE) and Southern blot with probes specific for the terminal C, I, L, and M fragments from chromosomes I and II. (C) PFGE analysis for N-terminal truncation mutants of Rap1. All mutants were V5-tagged at their C-terminus.

To further investigate a possible role for the PI domain in end protection, we next fused Rap1_491-693 to the C-terminus of Poz1 and integrated this construct at the endogenous *poz1* locus in cells, which possess elongated telomeres due to the absence of Rap1 in the starting strain. The fusion protein is expressed at lower than wildtype, but higher than Rap1_440-693 levels (Figure S1). Over the course of 14 successive restreaks on agar plates (approximately 310 generations), telomere length decreased and stabilized at slightly shorter than wildtype length (Figure 2A). Interestingly, at the 5th restreak (approximately 110 generations), cells characterized by intermediate telomere length showed mild end fusions, whereas no fusion phenotype was observed after 14 restreaks, when telomeres are no longer elongated (Figure 2B, compare lanes 3 and 4). These results confirm the dispensability of the PI domain under certain conditions, but more importantly, suggest that telomere length directly affects the degree of end protection with longer telomeres being more likely to undergo fusions than shorter ones.

**Figure 2.**
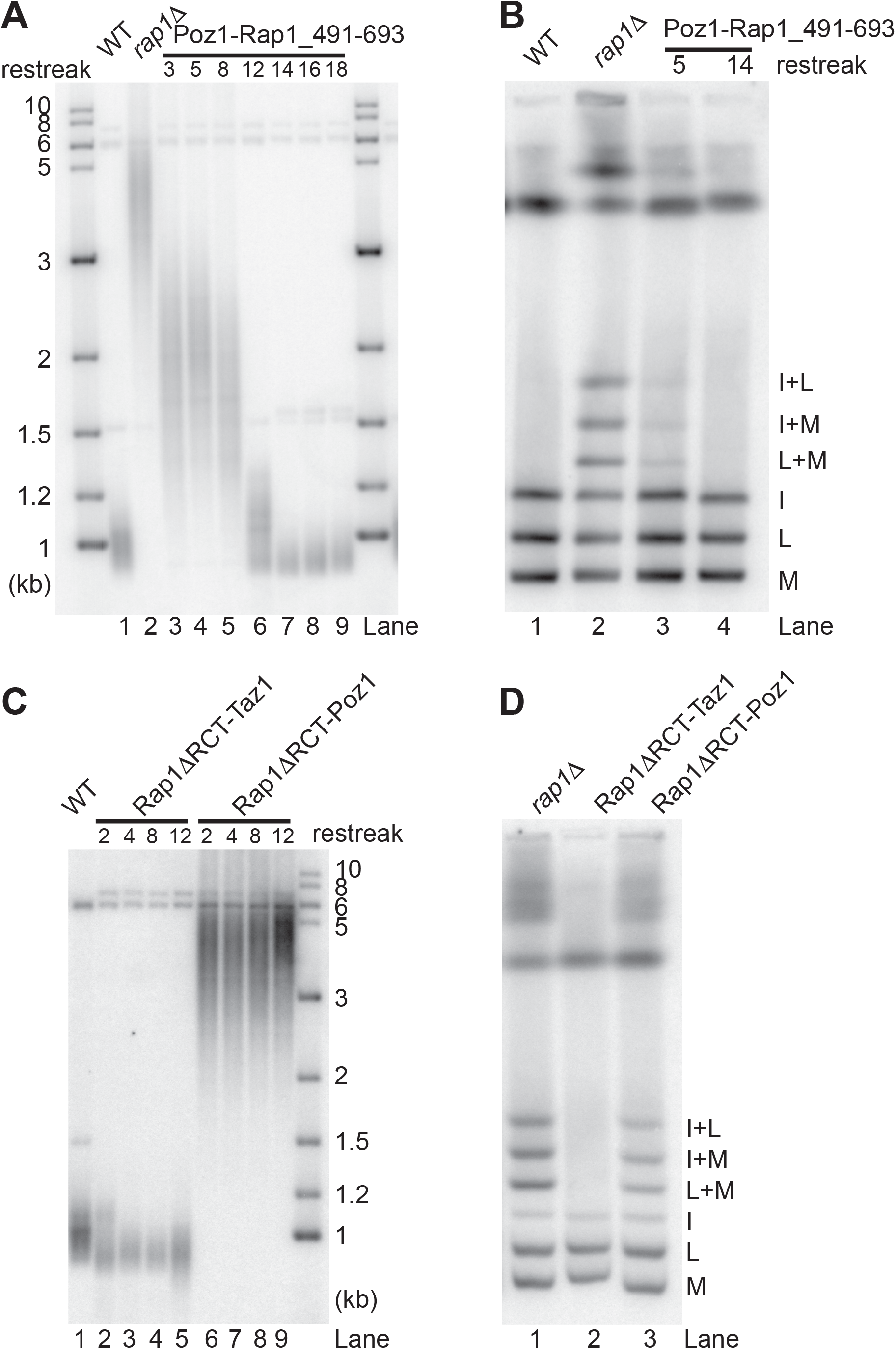
Covalent linkers can replace the PI and RCT domains of Rap1. (A) Telomere length analysis of cells from different sequential restreaks following introduction of Poz1-V5-Rap1_491-693 into *rap1Δ* cells. Telomere length was assessed by Southern blotting of EcoRI-digested genomic DNA probed with a telomere specific probe. (B) PFGE analysis of Poz1-V5-Rap1_491-693 after 5 and 14 restreaks. (C) Telomere length analysis of cells with Rap1ΔRCT fused to Taz1 or Poz1 with intervening V5 tag. The fusion protein was integrated at the endogenous *rap1* locus in the context of *taz1Δ* or *poz1Δ*, respectively. (D) PFGE analysis for Rap1ΔRCT-V5-Taz1 and Rap1ΔRCT-V5-Poz1.

To test whether the RCT domain itself is directly contributing to end protection beyond its function in the recruitment of Rap1 to telomeres, we fused Rap1ΔRCT to Taz1. This fusion protein rescued both the telomere length and end fusion phenotypes caused by the lack of the RCT domain (Figure 2C, lanes 2-5 and 2D, lane 2). In contrast fusing Rap1ΔRCT to Poz1 neither rescued telomere length nor end protection (Figure 2C, lanes 6-9 and 2D, lane 3). These results support that the PI and RCT domains are not directly involved in the protective role of Rap1. They are needed for the recruitment and stabilization of the protein but can each be replaced by covalent linkers to provide tethers to Poz1 and Taz1, respectively.

### Mutagenesis screen identifies residues important for end protection

As the results described above pointed to a function of the region between PI and RCT domains in end protection, we generated a library of randomly mutagenized Rap1_440-693 fragments and introduced them into *rap1Δ* cells (Figure 3A). The average mutation rate was 0.785% per nucleotide position as determined by Illumina sequencing of the library prior to introduction into *S. pombe*. We reasoned that cells harboring Rap1 mutations that impair end protection would be delayed in resuming cell division following G1 arrest, as chromosome end fusions occurring during G1 arrest will delay or prevent re-entry into the cell cycle. Hence, point mutations in Rap1 that affect its end protection function are expected to decrease in abundance over multiple rounds of arrest and return to growth (Figure 3A). After each round, cells were collected to assess telomere length and the occurrence of end fusions. The average telomere length and incidence of end fusions declined over the course of five rounds of arrest and return to growth (Figure S2).

**Figure 3.**
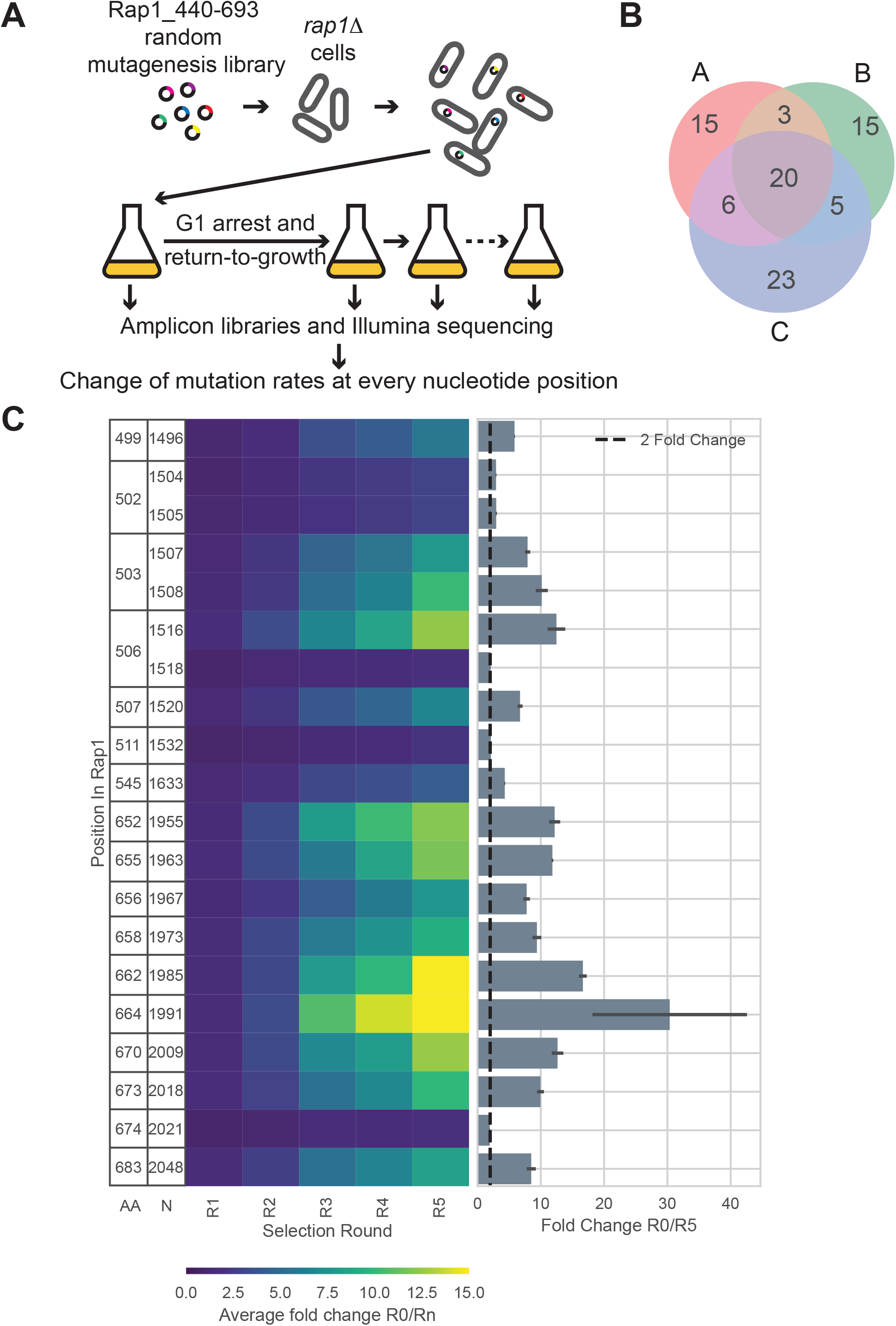
Random mutagenesis screen and Illumina sequencing identify important amino acid positions. (A) Schematic of the mutagenesis screen. A random mutagenesis plasmid library of Rap1_440-693 under the endogenous *rap1* promoter was introduced into *rap1Δ* cells. The cultures were subjected to five rounds of arrest and return to growth. Amplicon libraries covering the *rap1* fragments were generated and Illumina sequenced. Change of mutation rates at every nucleotide position were calculated. (B) Venn diagram of three biological replicates showing the positions where the non-native nucleotides consistently decrease from one round to the next. (C) Left panel: heat map showing the average changes in non-native nucleotides after each round of arrest and return to growth. Numbers represent the amino acid position in the full length protein (AA, first column) and the nucleotide position in cDNA (N, second column). Columns R1 to R5 represent rounds of selection. The color of the cells represents the average fold change in non-native nucleotides compared to the initial time point R0. Right panel: bar graph depicting average fold change for R5 over R0. Error bars show standard deviation. A dashed line denotes the 2-fold cutoff.

We then utilized Illumina sequencing to assess changes of mutation frequencies at each nucleotide position over time. An 877bp fragment including Rap1_440-693a.a. and short flanking regions was amplified from each round of the time course performed in triplicate. The mutation frequency profiles were similar before and approximately 12 generations after transformation into *S*.*pombe* confirming the preservation of library complexity upon transformation (Figure S3). After five rounds of G1 arrest followed by return to growth, the three replicates displayed striking overlap in which nucleotide positions showed a decrease in nucleotide substitution and which ones did not (Figure S4A). Confirming that the observed changes were indeed the result of selection at the protein level during growth in culture, the incidence of the mutant nucleotides decreasing over time was limited to codon positions 1 and 2, while position 3 remained mostly unchanged (Figure S4B).

Applying a filter to identify positions where non-native nucleotides consistently decreased each round in each of the replicates over the course of five rounds, we identified 20 nucleotide positions (Figure 3B), corresponding to 17 amino acids (Figure 3C). For three amino acids two codon positions were affected (Figure 3C and Table S1). In all cases the nucleotide positions under selection affected the encoded amino acids. After five rounds of arrest and return to growth, all except one position showed a greater than 2-fold change in favor of the native nucleotide (Figure 3C). In aggregate, these results confirmed the robustness of the assay and suggested that the identified amino acid positions must be important for end protection.

### Characterization of the identified amino acid positions

Of the positions identified in this screen, 10 clustered within the RCT domain (between amino acid 639 and 693) and 7 were found between position 491 and 638 (Figure 4A). No positions within the PI domain were under selection, further confirming that this region is dispensable for the protection from end joining. In contrast, mutations in the RCT domain are expected to disrupt the recruitment of Rap1 to telomeres. Indeed, several positions under selection make contact with Taz1 in the nuclear magnetic resonance structure (Chen *et al*., 2011), and we considered these positions as validation of the screen and did not further investigate the effect of the individual mutations in this region. Alignment of *S. pombe* Rap1 with two other fission yeast species, *S. octosporus* and *S*.*cryophilus*, showed that 5 of the 7 amino acids in the 491 to 638 region are identical in all three species and one represents a conserved valine to isoleucine change (Figure 4A). Interestingly, 6 of the 7 amino acids are within a 13 amino acid stretch (499-511) which at the time of the analysis was uncharacterized and we referred to as the protection patch or p-patch for short.

**Figure 4.**
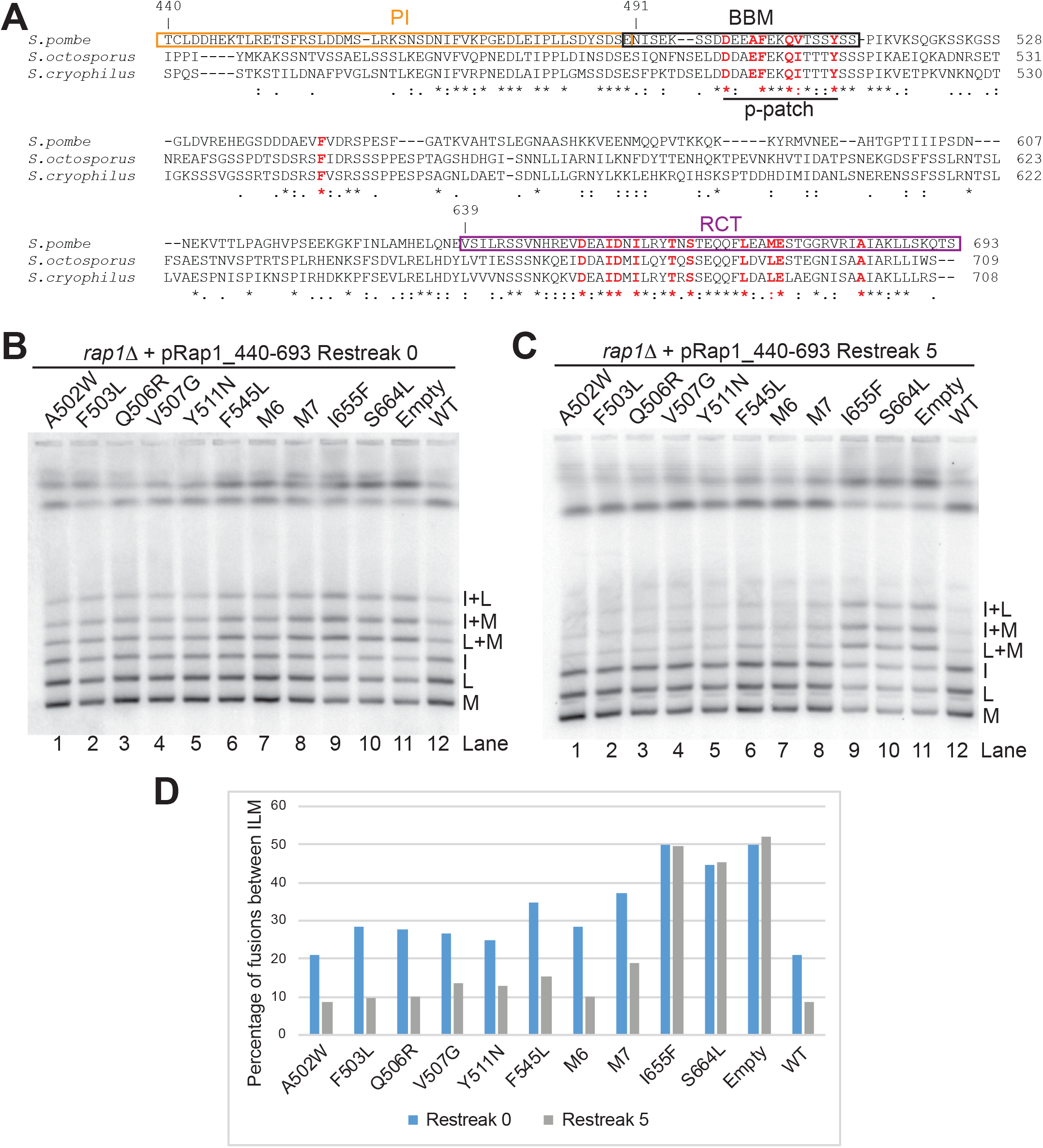
Important amino acids for telomere protection in Rap1_440-693. (A) Clustal Omega (1.2.4) alignment of Rap1 protein sequences from *S. pombe, S. octosporus* and *S. cryophilus* (Sievers *et al*, 2011). Amino acids identified in the mutagenesis screen are marked in red. Amino acids 499 to 511 harbored 6 of the selections and the region was named the p-patch. The previously described PI, BBM and RCT domains are shown for reference. (B) PFGE analysis of cells containing plasmids of Rap1_440-693 with mutations of the amino acids identified in (A) in the context of *rap1Δ*. Cells were subjected to G1 arrest after recovered from transformation (Restreak 0). (C) PFGE analysis of cells harboring the same mutations after 5 restreaks on plates (approximately 110 generations). (D) Quantification of (B) and (C) using Fiji. The y-axis shows the percentage of intensity represented by the three fusion bands (I+L, I+M and L+M) relative to the total intensity of the fused and unfused signals. WT: wildtype Rap1_440-693.

To confirm the importance of the identified positions, we created plasmids carrying the mutations A502W, F503L, Q506R, V507G, Y511N, and F545L, and introduced them into *rap1Δ* cells to assess whether protection from end fusions was compromised (Figure 4B and 4C). In addition, a compound mutant containing the 6 amino acids located within the p-patch was generated and named M6. Finally, all 7 positions identified to be under selection outside the RCT domain were combined in one construct (M7). As controls, we selected two amino acid changes in the RCT domain, I655F and S664L. I655 is part of the hydrophobic groove that interacts with Taz1 (Chen *et al*., 2011) and mutations of S664 were selected against most strongly of all the identified positions (Figure 3C). After recovery from transformation, cells were subjected to G1 arrest (Restreak 0) and analyzed by pulsed-field gel electrophoresis (PFGE) (Figure 4B). The two mutations in the RCT domain, I655F and S664L, behaved as null alleles showing similar levels of fusions as the empty vector control (Figure 4B, lanes 9-11 and 4D). The mutations in the p-patch and F545 displayed intermediate level of fusions (Figure 4B, lanes 1-8 and 4D). Interestingly, the wildtype C-terminal fragment of Rap1 also displayed fusions, albeit at lower levels than the point mutants (Figure 4B, lane 12 and 4D). These results confirmed that changes in Illumina sequencing read counts attributed to specific nucleotide changes are indeed a reflection of growth differences caused by varying levels of end fusions in the competition experiment.

To assess whether telomere length affected the incidence of end fusions, the cells carrying the 440-693 WT or mutant fragments were restreaked five times on plates (approximately 110 generations). In comparison to the cells analyzed immediately after transformation (Restreak 0), fusions were reduced by more than two-fold for the WT fragment, whereas fusions remained high in the RCT mutants I655F and S664L and the empty vector (Figure 4C, lanes 9-11 and 4D). Interestingly, all individual and the compound mutants in the p-patch and F545 showed decreased levels of fusions in Restreak 5 compared to Restreak 0 (Figure 4C, lanes 1-8 and 4D). Examination of telomere length revealed that Restreak 0 samples had long and heterogenous telomeres similar to *rap1Δ* cells (Figure S5A). In contrast, over the course of five restreaks telomeres had shortened substantially for the strains harboring WT, p-patch and F545 mutants, but not for the RCT mutants or empty vector (Figure S5B). These correlations in telomere length and incidence of end fusions when end protection is compromised strongly support that telomere length affects protection in a manner that makes longer telomeres more susceptible to fusions than shorter ones.

### Telomere length affects end protection

To further test the telomere length dependence of end protection, we integrated the point mutations in the p-patch and F545 in the context of full length *rap1* at the endogenous locus. In contrast to the previous experiment, cells now received the mutant alleles in the context of wildtype telomere length. Under these conditions, none of the point mutants displayed chromosome end fusions upon G1 arrest (Figure 5A). Even when full length Rap1 was replaced with the C-terminal fragment, the mutations did not result in compromised end protection (Figure 5B). Consistent with previous results that only the Poz1 and Taz1 interaction domains are required for telomere length maintenance (Pan *et al*., 2015), all mutants maintained wildtype telomere length in the context of full length and the C-terminal fragment (Figure S6A). To further test whether longer telomeres place an additional burden on end protection, we now triggered telomere lengthening by deleting *poz1*^*+*^ (Figure S6B). Strikingly, as telomeres elongated, fusions were readily detected in the p-patch (M6) and F545L mutants, but not in the presence of wildtype Rap1 (Figure 5C, lanes 2-4 and 5D). This result cannot be explained by differences in protein levels as the mutants were expressed at the same or slightly higher levels than WT Rap1 (Figure S7). When only the C-terminal fragment of Rap1 (440-693) was expressed, even the wildtype sequence was unable to fully protect elongated telomeres (Figure 5C, lane 5). Moreover, combining expression of only the C-terminal fragment with the point mutations had an additive effect on the level of chromosome end fusions (compare lanes 5 with 6 and 7). This effect was not due to differences in expression level (Figure S7B). No end fusions were observed when the gene encoding ligase IV was deleted, demonstrating that these are indeed mediated by the classical NHEJ pathway (Figure S8).

**Figure 5.**
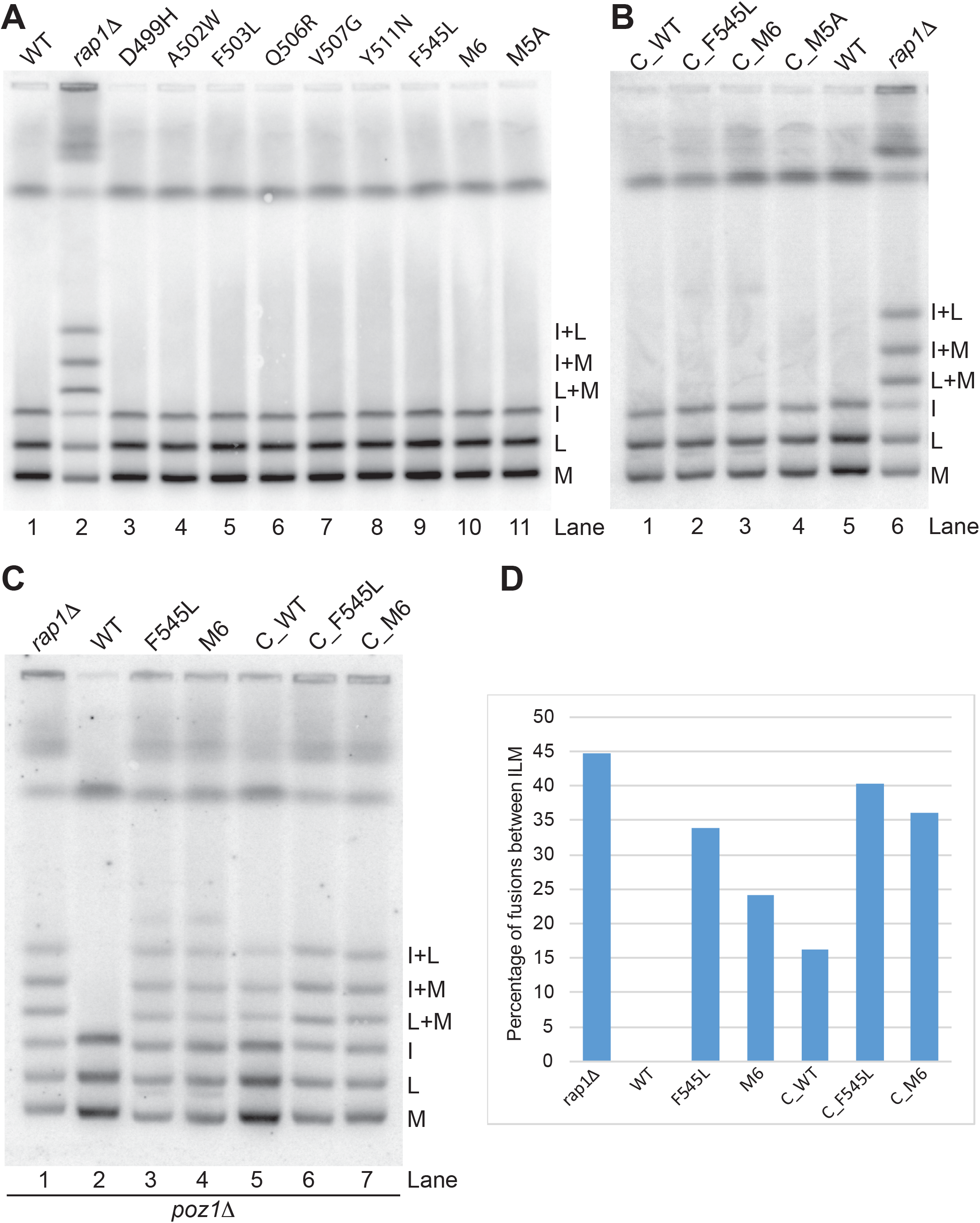
Rap1 mutants fail to protect long telomeres. (A) PFGE analysis of N-terminally V5-tagged full length Rap1 mutants. All mutants were integrated at the endogenous *rap1* locus. M5A is the mutation of all 5 non-alanine amino acids under selection in the p-patch to alanine. (B) PFGE analysis of N-terminally V5-tagged *rap1*_440-693 mutants integrated at *rap1* endogenous locus. (C) PFGE analysis of *rap1* mutants in *poz1Δ* background. (D) Quantification of (C) by Fiji as in Figure 4D.

To further test whether the p-patch and F545 are specifically required for the protection of long telomeres, we created four internal deletions within the region between the PI and RCT domains (Figure 6A). Western blot analysis showed that the expression levels for WT and all deletion mutants were within 2.5-fold of each other with WT and Δ593-638 at the lower end of the range (Figure 6B). The three non-overlapping deletion mutants exhibited wildtype telomere length (Figure 6C, lanes 4-6) and no end fusion phenotype (Figure 6D, lanes 3-5). When the entire 491-638 region was deleted, we observed slight telomere elongation (Figure 6C, lane 3) and mild fusions (Figure 6D, lane 6), but only in the presence of the V5 epitope tag (compare Figure 6C, lane 3 and 7; Figure 6D lane 6 and 7). These results demonstrate that the entire region between the PI and RCT domains is dispensable for end protection if telomeres are wildtype in length.

**Figure 6.**
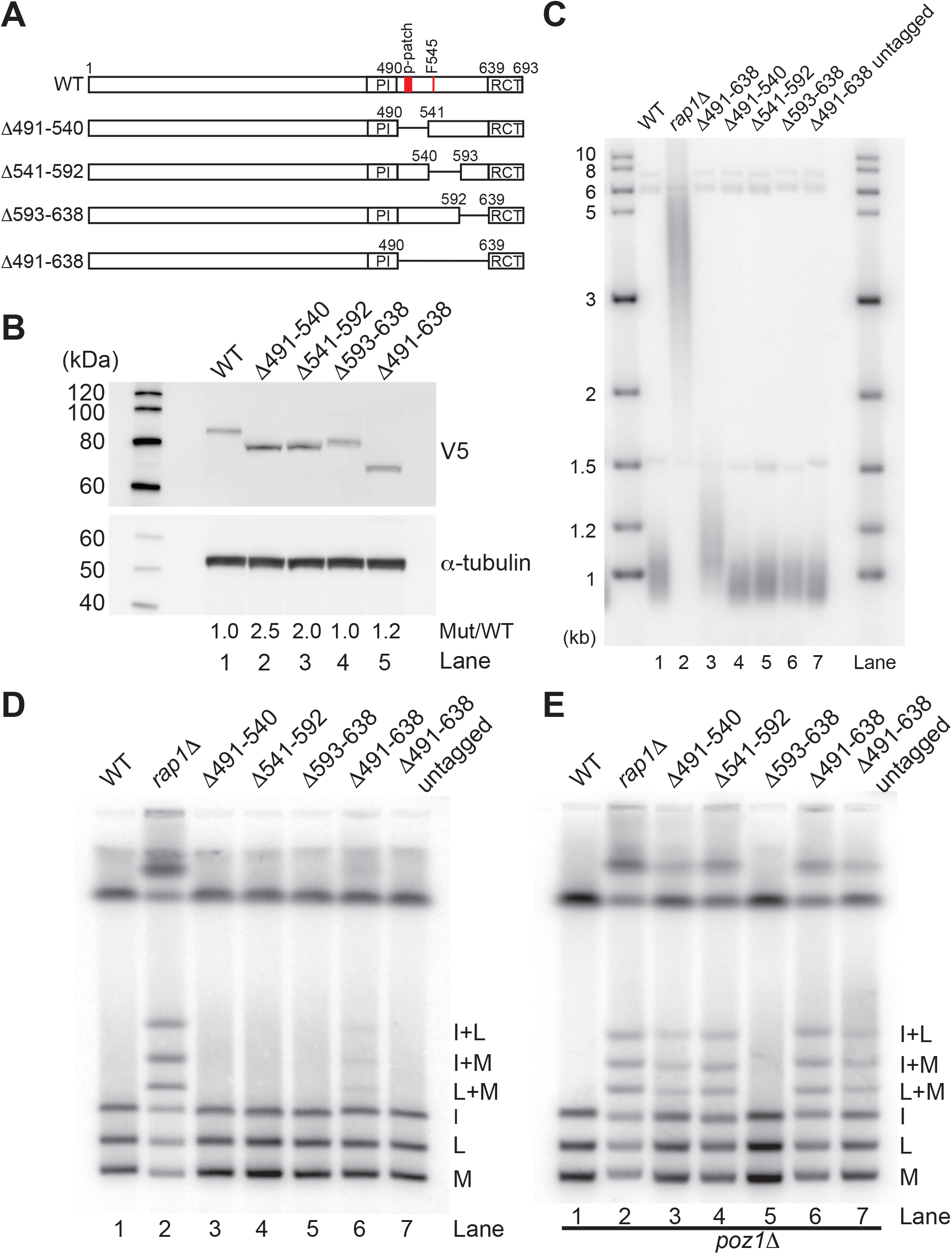
The region between the PI and RCT domain of Rap1 is not necessary for end protection when telomere length is wildtype. (A) Schematics of the internal deletion mutants between PI and RCT domains. (B) Western blot analysis of N-terminally V5-tagged truncation mutants using anti-V5 antibody. Anti-α-tubulin was used as loading control. Quantification of Rap1 mutant protein levels relative to WT normalized to α-tubulin is shown beneath the gel. (C) Telomere length analysis of the mutants by Southern blot with telomere specific probe. All mutants are V5-tagged except for Δ491-638 untagged (lane 7). (D) PFGE analysis of the same strains. (E) PFGE analysis in *poz1*Δ background.

In contrast, when telomeres were elongated by deletion of *poz1*, Rap1Δ491-540, lacking the p-patch, and Rap1Δ541-592, where F545 resides, both displayed end fusion phenotypes (Figure 6E, lanes 3 and 4); whereas Rap1Δ593-638, where no positions important for end protection were identified in our screen, showed full protection even in the presence of very long telomeres (lane 5). These results further support a specific function for the p-patch and F545 in the protection of long telomeres.

### Nuclear envelope attachment mediates the protection of long telomeres

Following our characterization of the p-patch mutants, a biochemical and structural analyses of the Rap1-Bqt4 interaction was published (Hu *et al*, 2019) and revealed that the p-patch overlaps with the Bqt4 binding motif (BBM) of Rap1. The Rap1-Bqt4 interaction tethers telomeres to the nuclear envelope (NE), and is required during meiosis for telomere clustering, meiotic recombination and spore formation (Chikashige *et al*, 2009). In contrast, this attachment is largely dispensable for the maintenance of telomere length and the silencing of subtelomeric genes during mitotic growth. Considering the overlap of p-patch and BBM, we wondered whether the nuclear envelope attachment of telomeres was important for the protection of long telomeres during vegetative growth. Telomere localization within the nucleus is commonly examined by defining three concentric nuclear zones of equal surface area ((Hediger *et al*, 2002), Figure 7A). Pot1-GFP was used to visualize telomeres and Nup44-mCherry to label the NE. Consistent with previous results that loss of *bqt4* releases telomeres from the NE (Chikashige *et al*., 2009; Maestroni *et al*, 2020), a significant decrease in the fraction of telomeres in zone I (nuclear periphery) was observed in *bqt4* deletion cells (Figure 7B). Localization of telomeres in zone I was diminished to the same extent in the Rap1 M6 mutant (Figure 7B). These results were confirmed when telomeres were visualized by Taz1-GFP (Figure S9A). In summary, we conclude that the p-patch mutants impair NE attachment presumably by disrupting the Rap1-Bqt4 interaction. Interestingly, the F545L mutant appears to affect end protection by other, as of yet uncharacterized, means. Although F545L caused a specific loss of end protection for long telomeres similar to the mutations in the p-patch, this mutant is not part of the Rap1-Bqt4 interaction interface (Hu *et al*., 2019) and did not show a decrease in telomere localization to the nuclear periphery (Figure 7B).

**Figure 7.**
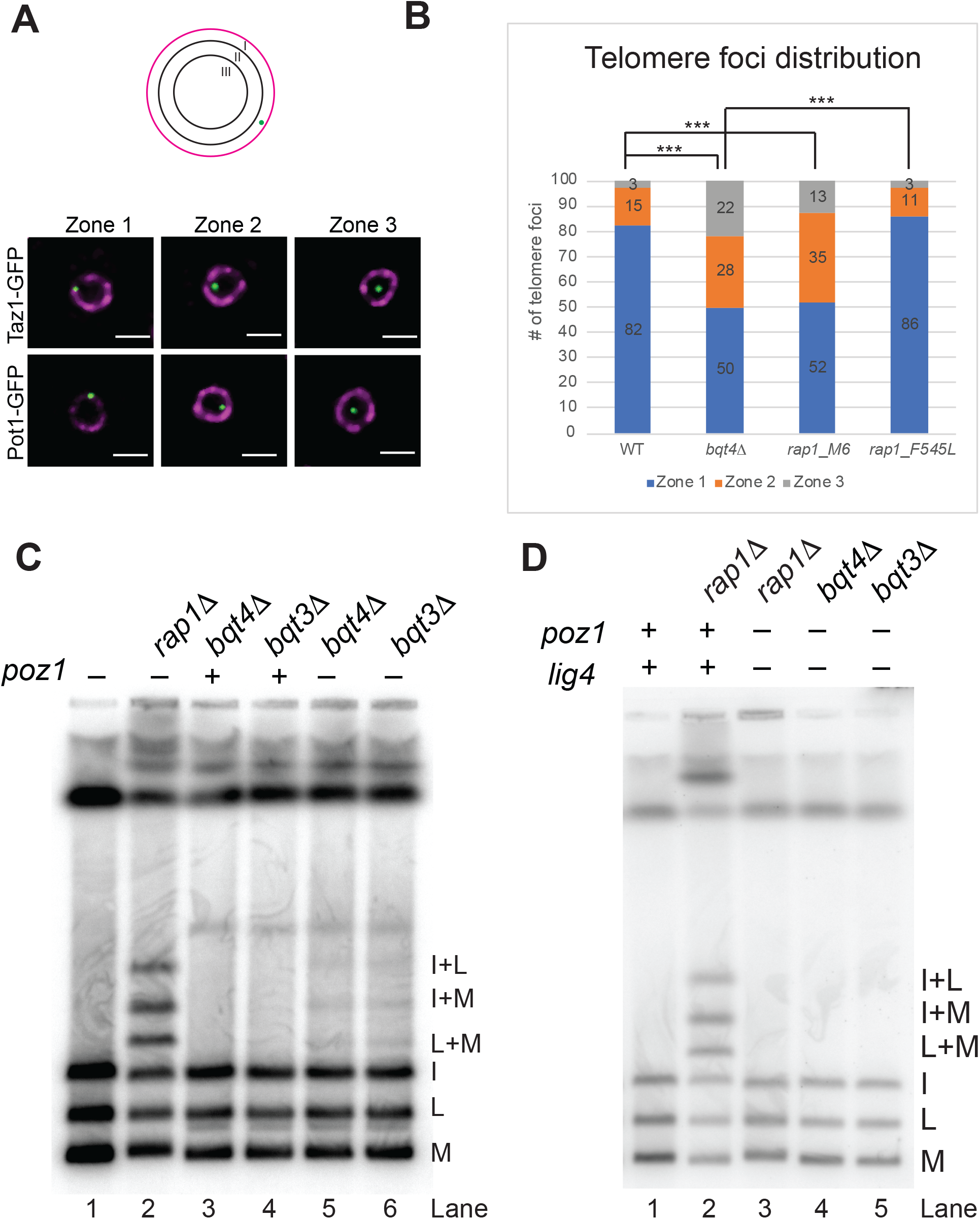
Nuclear envelope attachment mediates end protection of long telomeres. (A) Schematic of scoring matrix. Each nucleus was divided into three concentric zones of equal area with zone I corresponding to the nuclear periphery. Distance of the GFP spot to NE and diameter of the nucleus were measured for the calculation. Representative images of cells expressing Nup44-mCherry (magenta) and Taz1-GFP or Pot1-GFP (green), respectively. The z-plane showing the brightest GFP spot is shown and was used for quantification. Scale bar: 2μm. (B) Stacking bar chart showing telomere distribution (labeled by Pot1-GFP) in each of the three zones. 100 telomere foci were quantified per sample. Chi square tests were performed by comparing the *rap1* mutants to WT or *bqt4Δ*. Significance was indicated. ***: p < 0.001, **: 0.001 <= p < 0.01, *: 0.01 <= p < 0.05. (C) PFGE analysis of *bqt4*Δ and *bqt3*Δ cells in the presence or absence of *poz1*. (D) End fusions in *bqt4*Δ*poz1Δ* and *bqt3*Δ*poz1Δ* are dependent on the presence of ligase IV.

To further test whether protection of long telomeres specifically requires NE attachment, PFGE was performed with genomic DNA from *bqt4*Δ and *bqt3*Δ cells. Bqt3 also localizes in the inner nuclear membrane and interacts with and protects Bqt4 (Chikashige *et al*., 2009). No end fusions were observed in *bqt4*Δ or *bqt3*Δ cells when telomere length was normal (Figure 7C, lanes 3 and 4). However, when telomeres were elongated by loss of Poz1 (Figure S9B), fusions were detected in both *bqt4*Δ and *bqt3*Δ cells (Figure 7C, lanes 5 and 6). The dependence of the fusions on ligase IV confirmed that they are mediated by NHEJ (Figure 7D). In aggregate, these results demonstrate an important role for NE attachment in specifically protecting long telomeres. In addition, another mechanism is likely involved as the mutation of F545 compromises the protection of long telomeres without affecting NE attachment. These findings are consistent with the notion that multiple and partially redundant pathways are involved in mediating the important function of safeguarding chromosome ends.

## Discussion

Rap1 plays key roles in limiting telomere elongation and protecting chromosome ends from fusions. While its role in regulating telomere length is mainly based on it being part of a protein bridge that tethers the single- and double-stranded parts of the telomere (Kim *et al*, 2017; Pan *et al*., 2015), its contributions to preventing chromosome end fusions are more complex. Using molecular genetic approaches, we have now identified several amino acid positions within Rap1 that are specifically important for the protection of long telomeres. Our results reveal that the requirements for end protection vary depending on the length of a telomere and that one of the mechanisms by which Rap1 protects chromosome ends is by limiting telomere elongation itself. Already elongated telomeres are protected by tethering them to the nuclear periphery, a function that is not required for the protection of shorter telomeres.

Examining end protection in cells with wildtype telomere length revealed a surprising flexibility in terms of the required structural elements. Deletions that in aggregate cover the entire protein (Δ1-439, Δ440-490, Δ491-540, Δ541-592, Δ593-638, Δ639-693) showed that any specific region can be deleted or replaced with a simple linker while maintaining the protective function of the protein (Figures 1, 2 and 6). These results demonstrate that any element within Rap1 that is involved in end protection functions in a redundant manner. Considering the catastrophic consequences that follow failure to protect chromosome ends from undergoing fusions, redundancy is perhaps not surprising. Similarly, Rap1 in budding yeast inhibits NHEJ through multiple mechanisms(Marcand *et al*, 2008). Here, Rap1 binds telomeric DNA directly and its RCT domain engages Rif2 and Sir4 to inhibit end fusions in two parallel pathways. In addition, the central region of Rap1, which harbors the DNA binding domain but excludes the BRCT and RCT domains, has an independent function in end protection that is only revealed when *rif2* and *sir4* are deleted. In mammalian cells, end protection by Trf2 masks any role of Rap1 under some conditions (Sfeir *et al*., 2010). Protection of chromosome ends by Rap1 is only observed when Trf2 is absent (Sarthy *et al*., 2009) or when cells enter replicative senescence (Lototska *et al*., 2020).

In dissecting the role of Rap1 in end protection, we realized that telomere length profoundly affects end protection. However, in contrast to the long-established view that the shortest telomeres are most at risk of triggering a DNA damage response, we found that long telomeres are also highly vulnerable to end fusions. Rap1 mutants that fully protect wildtype telomeres, fail to prevent fusions of long telomeres. This is observed independently of whether telomere elongation was induced by the deletion of *poz1* (Figure 5C and 6E), or by using a genetic background with initially long telomeres that shortened over time (Figure 2B, 4B and 4C).

One explanation for the telomere length dependence of end protection is that elongated telomeres require more telomeric proteins to coat all telomeric repeats. While we cannot exclude the possibility that limiting protein levels provide a partial explanation for our observations, two reasons argue against insufficient Rap1 being responsible for the loss of end protection. Firstly, Rap1 protein is expressed in large excess over what is needed to protect telomeres as demonstrated by its ability to fully protect 10-fold elongated telomeres (Figure 5C). Secondly, the protein levels of several mutants were higher than WT, yet they failed to protect against fusion of elongated telomeres (Figure 5C and 6E), even when telomeres are only slightly longer than wildtype (Figure 6D). Most importantly, Rap1ΔPI and Rap1Δ593-638, two mutants with non-overlapping deletions, fully inhibit end fusions in the presence of long telomeres, even though Rap1Δ593-638 was expressed at the lowest level of all internal deletion mutants (Figure 1B, 6B and 6E). In contrast, the mutations at the positions identified in the screen failed to protect elongated telomeres, strongly indicating a functional significance of the p-patch and F545 in specifically protecting long telomeres.

Mechanistically, the mutations within the p-patch disrupt the tethering of telomeres to the nuclear periphery. Consistent with the loss of this tether rendering long telomeres more vulnerable to undergoing fusions, deletion of *bqt3* or *bqt4* phenocopies this sensitivity. Furthermore, TERRA expression and telomere recombination are increased when telomeres detach from the nuclear envelop (Maestroni *et al*., 2020). Similarly in human cells, telomeres localize to the nuclear periphery during post-mitotic nuclear assembly (Crabbe *et al*, 2012)and nuclear envelop tethering has been shown to inhibit the formation of ALT-associated PML bodies and by inference telomere recombination (Yang *et al*, 2021). Another recent study found that telomeres experiencing replication stress due to the presence of the Pot1ΔOB mutant show relocalization of telomeres to the nuclear periphery possibly to preserve telomere integrity (Pinzaru *et al*, 2020). Notably, the presence of Pot1ΔOB in human cells causes extreme telomere lengthening (Loayza & De Lange, 2003) indicating that the presence of a protective mechanism for long telomeres in the nuclear periphery may well be conserved between fission yeast and human cells. In contrast, the localization of telomeres to the nuclear periphery in budding yeast has been proposed to promote genome integrity by stimulating DSB repair of subtelomeric breaks (Schober *et al*, 2009; Therizols *et al*, 2006). However, differences between long and short telomeres were not examined in these studies. It is clear that tethering of telomeres to the nuclear envelop puts them in a privileged compartment, although the molecular consequences may vary among species and circumstances.

Telomere length is highly heterogenous across evolution, but each species maintains telomeres within a defined size range. This is true even in species with constitutively active telomerase, where shortening of telomeres cannot serve as a tumor suppressive mechanism. It is conceptually easy to see that cells maintain a lower limit of telomere length to preserve the mark that distinguishes a chromosome end from the DNA ends formed by a DNA double-strand break. As telomeres erode past a critical length, a DNA damage response may result in chromosome end fusions which, in the event of continued cell division, causes genomic rearrangements or mitotic catastrophe (Armanios, 2009; Artandi & DePinho, 2010; De Lange, 2005). In contrast, why cells would maintain an upper limit of telomere length is less clear, even though it has been known for some time that long telomeres pose challenges. Telomeres are fragile sites susceptible to replication fork stalling that require the aid of shelterin components for efficient replication (Miller *et al*, 2006; Sfeir *et al*, 2009). Their G-rich nature also render telomeres more susceptibility to oxidative damage and longer telomeres are more sensitive to high levels of reactive oxygen species (Rubio *et al*, 2004). Now, our results show that longer telomeres require specific protection against fusions, revealing another challenge for cells with long telomeres to maintain genome integrity. It is thus not surprising that cells have active mechanisms to shorten hyper-elongated telomeres such as telomere rapid deletion events (TRD) or telomere trimming (Li & Lustig, 1996; Pickett *et al*, 2009; Wang *et al*, 2004).

Interestingly, several recent studies have reported an association between long telomeres and an increased risk for various types of cancer (Julin *et al*, 2015; Lan *et al*, 2009; Lynch *et al*, 2013; Machiela *et al*, 2017; Machiela *et al*, 2015; Machiela *et al*, 2016; Ojha *et al*, 2016; Pellatt *et al*, 2013; Rode *et al*, 2016). The authors reason that long telomeres provide cells with additional time to accumulate driver mutations while proliferating before short telomeres trigger cell cycle arrest. Our data now suggests an additional, but not mutually exclusive explanation: long telomeres may themselves increase genomic instability and thereby drive tumorigenesis.

## Materials and Methods

### Strains and Constructs

All strains used in this study are listed in Table S2. Genomic integrations were generated by one-step gene replacement (Bahler *et al*, 1998). Plasmids used in this study are listed in Table S3 and were transformed into *S. pombe* by electroporation. Plasmid pLP150 contained *rap1* 440-693 region flanked by 686 base pairs (bp) of the *rap1* endogenous promoter region plus an ATG start codon on the 5’ side and 375 bp of endogenous downstream terminator sequence inserted into pDBlet vector (Brun *et al*, 1995) and was isolated from the mutagenesis library described below. Plasmids pLP161-171 were generated by site-directed mutagenesis using pLP150 as template.

### Telomere length and end fusion analysis

Genomic DNA preparation, telomere length analysis and pulsed-field gel electrophoresis were carried out following the same procedure as described (Pan *et al*., 2015). Fiji (Schindelin *et al*, 2012) was used for the quantification of Southern blots by comparing combined signal intensities of I+L, I+M and L+M bands to the total intensity of the I, L, M bands and the three fusion bands.

### Denatured protein extract and western blotting

Extract preparation and western blot analysis was carried out as described (Pan *et al*., 2015). Primary antibodies used were mouse anti-V5 (Thermo Fisher Scientific, R960-25) and mouse anti-α-tubulin (Sigma-Aldrich, T5168). HRP-conjugated goat anti-mouse IgG (H+L) (Thermo Fisher Scientific, 31430) was used as secondary antibody.

### Mutagenesis library construction and screen

Oligos and DNA templates used for the generation of the mutagenesis library and PCR products for sequencing are listed in Table S4. Type IIS restriction enzymes were used to avoid introduction of non-native sequences between the coding region and the promoter and terminator sequences, respectively. First, Rap1 upstream 686bp and downstream 375bp regions were amplified by fusion PCR placing two BspMI restriction sites in opposite orientations between the upstream and downstream sequences, followed by cloning into pDblet with SalI and XmaI restriction enzymes to generate pLP143. The resulting plasmid was then digested with BspMI and treated with recombinant shrimp alkaline phosphatase (rSAP) prior to ligation. Rap1 fragment 440 to 693 with added start and stop codons and flanking BmsBI sites was amplified and TOPO-cloned to pCR4-Blunt-TOPO vector (Invitrogen) and sequence verified (pLP142). Random mutagenesis PCR was carried out with GeneMorph II Random Mutagenesis Kit (Agilent) following the manufacturer’s instructions with the following specifics: 5.95ng of pLP142 equivalent to 1ng of target DNA was used as template and 30 cycles for PCR. The product was digested with BsmBI and inserted into plasmid pLP143 by ligation. The ligation product was transformed to XL10-gold ultracompetent cells. To preserve the complexity of the library, 12 transformations with 2µl of ligation reaction per transformation were performed and plated to 73 large (150mm) plates containing Luria Broth (LB) supplemented with 50 μg/ml carbenicilin. From each plate ∼900 colonies were collected by scraping with a spreader and washing off with LB. Cell pellets were collected by centrifugation at 6000xg for 15min and plasmid DNA was isolated using a QIAfilter Plasmid Giga Kit (Qiagen).

The mutagenesis library plasmids were transformed into PP1623A (*rap1::natMX*) cells by electroporation. Cells from 6 transformations with 1µg of DNA per transformation were pooled after electroporation and allowed to recover in 30ml YES for 3 hours at 32°C. Cells were collected by centrifugation and transferred into 200ml EMM for overnight culture. Cultures were then diluted to 5×10^5^ cells/ml in 400ml EMM and cultured to mid-log phase (0.5-1 x10^7^ cells/ml). Cells were spun down and washed twice with EMM minus nitrogen (EMM-N), then resuspended in 360ml EMM-N at 2 x10^6^ cells/ml, split into 3 flasks as triplicates (A, B, C) and arrested at 25°C for 48h. The remaining cells were harvested for genomic DNA isolation as R0 sample. Following arrest, 4 x10^8^ cells were resuspended in 100ml EMM for each replicate and recovered for 6 hours and then diluted to 5×10^5^ cells/ml in 400ml with EMM and cultured at 32°C overnight. When the cell density reached 0.5-1 × 10^7^ cells/ml, 2.5 x10^8^ cells were collected by centrifugation, washed twice with EMM-N and resuspended in 125ml EMM-N at 2 × 10^6^ cells/ml for a second round of arrest at 25°C for 48h. These arrest-recovery cycles were repeated 5 times. Each round, cells were harvested for PFGE analysis following arrest and after recovery to mid-log phase, cells were harvested for genomic DNA preparation.

### Illumina sequencing and data analysis

Genomic and plasmid DNA from R0 and three replicates of R1 to R5 was isolated as described (Pan *et al*., 2015). To amplify the mutated region for Illumina sequencing, PCR reactions (50 µl) containing 1 × Q5 reaction buffer (NEB), 200 µM dNTPs, 0.5 µM of Bloli6118 and Bloli6119, 1 U of Q5 Hot Start High-Fidelity DNA Polymerase (NEB) and 100ng genomic DNA from R0-R5 or 93pg of the original plasmid library isolated from bacteria were incubated at 98°C for 1min, followed by 18 cycles of 98°C for 10s, 56°C for 30s and 72°C for 30s with a final extension at 72°C for 2min. PCR products were gel purified with MinElute gel purification kit (Qiagen) and Illumina libraries were prepared with different indexes using Nextera XT Library Prep Kit (Illumina, FC-131-1096). Prior to sequencing, the libraries were pooled at equal molar ratios and spiked in at 3.55% into a pool of unrelated RNA seq libraries to increase complexity of the sequencing samples and facilitate cluster calling. The libraries were then sequenced using the Illumina HiSeq-2500 platform in rapid mode on a RapidSeq flowcell at 100bp single read length. The number of reads generated per library is listed in Table S5.

Sequences were trimmed using Trimmomatic version 0.32 (Bolger *et al*, 2014) as follows: The leading and trailing bases of each read were removed if the quality score was below 3. Reads were scanned 5’ to 3’ with 4-base window and were clipped when the average quality score dropped below 15. After window clipping and leading/trailing bases were trimmed, reads less than 36bp long were dropped. Reads were then aligned to the modified Rap1 sequence using the bwa mem aligner version 0.7.15-r1140 (Li & Durbin, 2009). The resulting alignments were filtered for reads with MAPQ scores greater than or equal to 2, sorted and then indexed using SAMtools version 1.3.1. Per-position base composition of each alignment was determined using the command line tool pysamstats (https://github.com/alimanfoo/pysamstats; https://github.com/pysam-developers/pysam) (Li *et al*, 2009). The resulting tab-separated files were analyzed using Python version 2.7.2, and the data analysis package pandas (McKinney, 2010). Positions in each replicate were selected if the fraction of non-canonical nucleotide decreased from each round of selection to the next. The intersection of these positions was taken across replicates.

### Fluorescence microscopy and quantification

Live G1 arrested cells (in EMM-N for 24h) were mounted on 8 well glass bottom slides (ibidi) coated with concanavalin A and analyzed at 25°C. Fluorescence microscopy images were obtained using a fluorescence spinning disc confocal microscope, VisiScope 5 Elements (Visitron Systems GmbH), which is based on a Ti-2E (Nikon) stand equipped with a spinning disc unit (CSU-W1, 50 µm pinhole, Yokogawa). The set-up was controlled by the VisiView software (Visitron Systems GmbH) and images were acquired with a 100 × oil immersion objective (100 × NA 1.49 lens, Apo SR TIRF) and a sCMOS camera (BSI, Photometrics). Taz1-GFP and Pot1-GFP were detected with 488nm excitation laser and ET460/50 emission filter (Chroma). Nup44-mCherry was detected with 561 nm excitation laser and ET570LP emission filter (Chroma). 3D stacks of images with 14 confocal z-planes at 0.32-µm increments were recorded for each sample. The exposure time was 200 ms for each channel with a live bin of 1.

Fiji was used for image processing and quantification of the telomere foci distribution. Sample identities were blinded for the person performing image acquisition and quantification. In cells with large and well-defined nuclear envelope signal, the focal plane with the brightest GFP signal was chosen for measurements. The diameter of the nuclear envelope and the distance of the GFP spot to the nearest point of the nuclear envelope (X = distance between center of the spot and center of the NE signal) were measured. GFP signals were allocated to one of the three concentric zones of equal surface area as described (Hediger *et al*., 2002). Zone I (X <0.184 × radius (r)) is considered the nuclear periphery. X between 0.184 and 0.422 times r is zone II and X >0.422 × r is zone III. For each sample, 100 telomere foci were quantified. Chi square tests were performed by comparing the mutants to WT and *bqt4Δ*, respectively.

## Data availability

The sequencing data is available at GSE190759. Scripts used in this study are available at https://github.com/baumannlab/Sp_Pan_rap1_select_detect

## Acknowledgements

We thank Yasushi Hiraoka and Sue Jaspersen for strains, the Molecular Biology and Media Preparation facilities at the Stowers Institute for Medical Research and the Media Lab, Microscopy and Genomics Core Facility at the Institute of Molecular Biology for excellent service and support, David V Ho for computational help, Kristi Jensen for comments on the manuscript and all members of the Baumann laboratory for helpful discussions. This work was supported in part by the Deutsche Forschungsgemeinschaft (DFG; German Research Foundation; project number 393547839 – SFB 1361, sub-project 17), the Max Planck Graduate Center with the Johannes Gutenberg-University Mainz, the Howard Hughes Medical Institute and the Stowers Institute for Medical Research. P.B. is an Alexander von Humboldt Professor at Johannes Gutenberg University.

## Author Contributions

L.P. and P.B. designed the experiments; L.P. performed the experiments except for acquiring the microscopy data (N.B). The analysis of the Illumina sequencing data from the mutagenesis screen was performed by D.T.; all authors analyzed the data and L.P. and P.B. prepared the manuscript.

---

**Table S1:** Rap1 nucleotide positions selected from the screen.

| Raw data position<sup>a</sup> | cDNA nucleotide position | WT nucleotide | Codon | Codon Position | Amino Acid<sup>b</sup> | Mutation tested |
| --- | --- | --- | --- | --- | --- | --- |
| 255 | 1496 | A | GAT | 2 | D499 | D499H |
| 263 | 1504 | G | GCT | 1 | A502 | A502W |
| 264 | 1505 | C | GCT | 2 | A502 | A502W |
| 266 | 1507 | T | TTT | 1 | F503 | F503L |
| 267 | 1508 | T | TTT | 2 | F503 | F503L |
| 275 | 1516 | C | CAA | 1 | Q506 | Q506R |
| 277 | 1518 | A | CAA | 3 | Q506 | Q506R |
| 279 | 1520 | T | GTG | 2 | V507 | V507G |
| 291 | 1532 | A | TAC | 2 | Y511 | Y511N |
| 392 | 1633 | T | TTT | 1 | F545 | F545L |
| 714 | 1955 | A | GAT | 2 | D652 | NA |
| 722 | 1963 | A | ATC | 1 | I655 | I655F |
| 726 | 1967 | A | GAC | 2 | D656 | NA |
| 732 | 1973 | T | ATT | 2 | I658 | NA |
| 744 | 1985 | C | ACA | 2 | T662 | NA |
| 750 | 1991 | C | TCA | 2 | S664 | S664L |
| 768 | 2009 | T | CTT | 2 | L670 | NA |
| 777 | 2018 | T | ATG | 2 | M673 | NA |
| 780 | 2021 | A | GAA | 2 | E674 | NA |
| 807 | 2048 | C | GCC | 2 | A683 | NA |

a. Raw data position number is based on the PCR fragment amplified from the library which was used for Illumina sequencing
b. Amino acid number based on the amino acid position in the full-length protein
NA: not tested

**Table S2.**
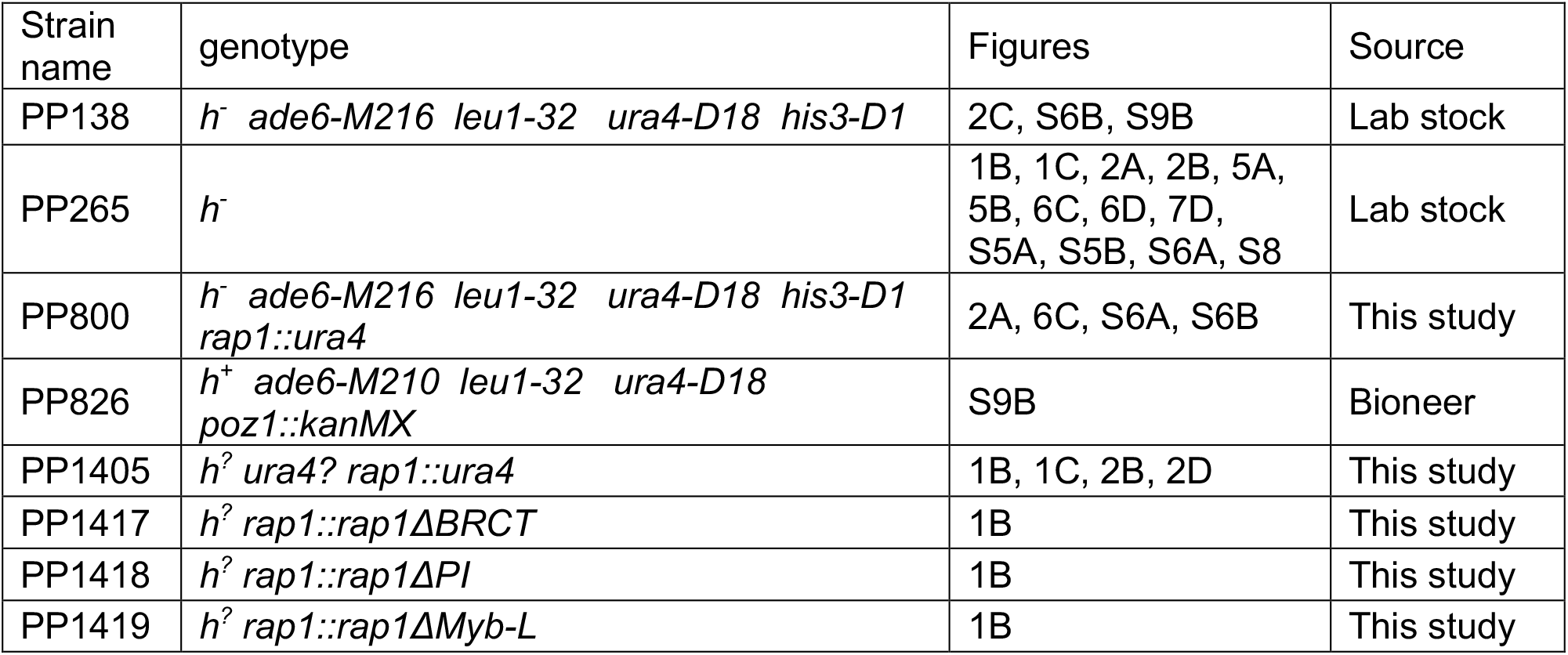
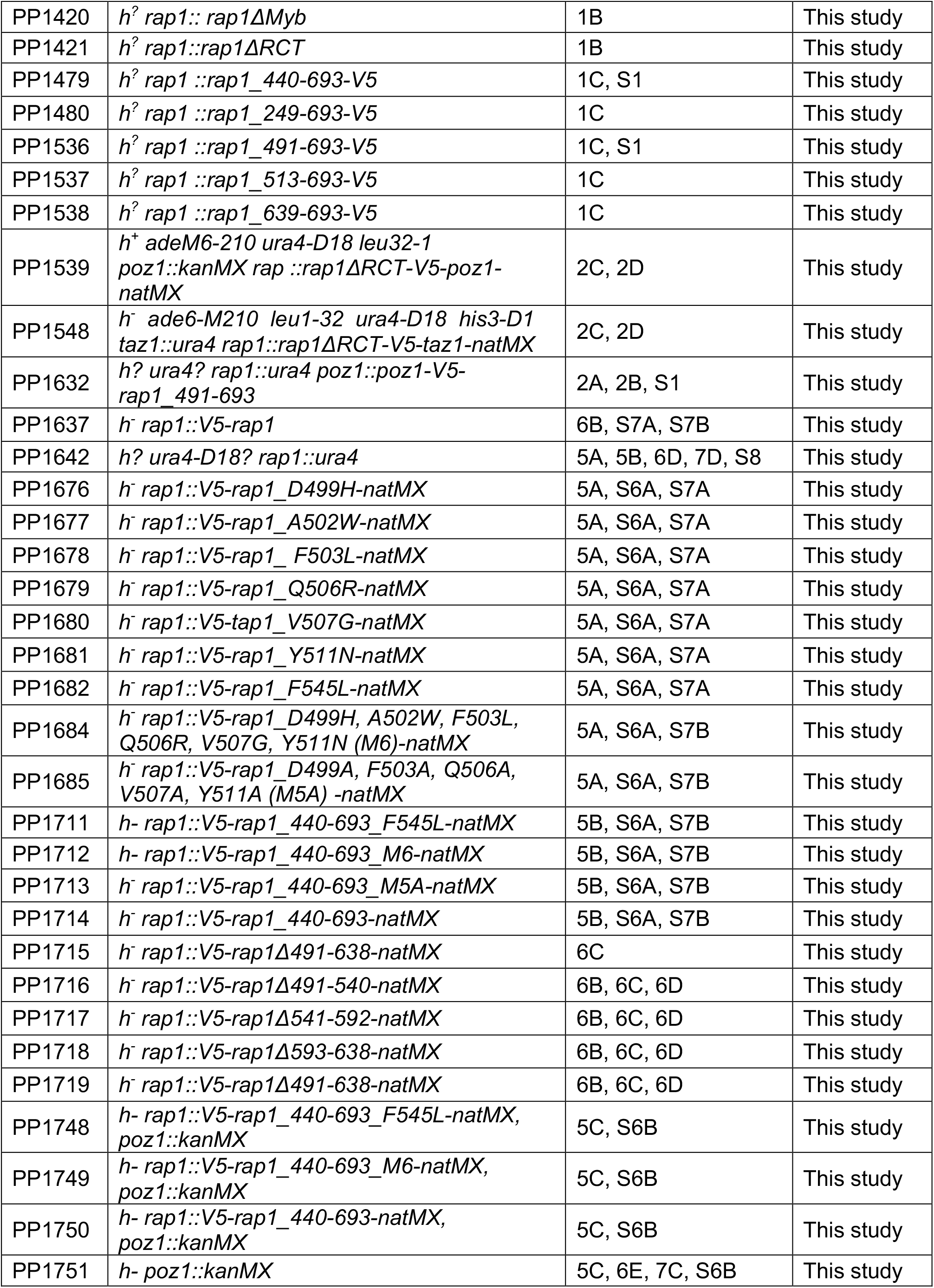
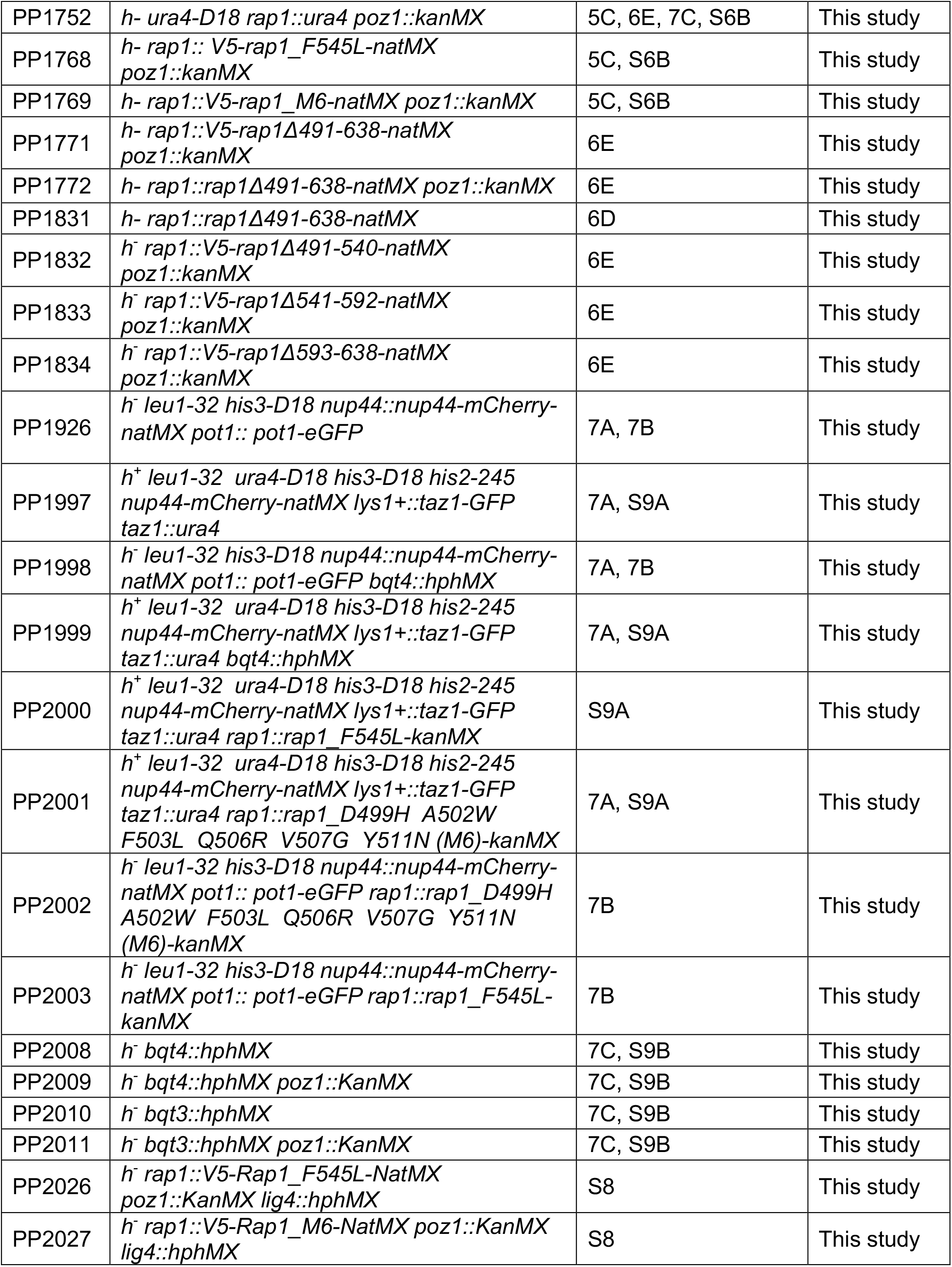
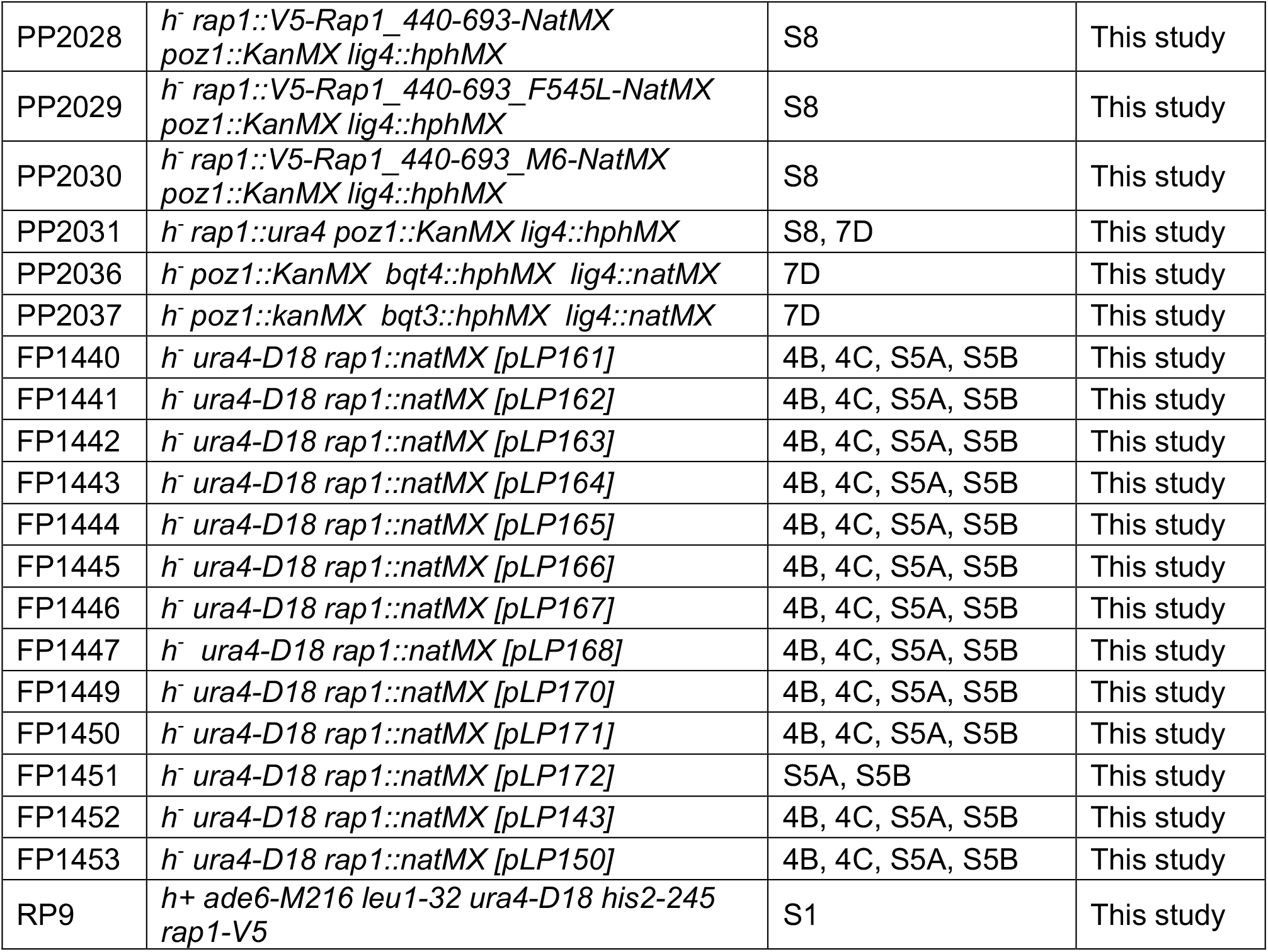
Strains used in this study.

**Table S3.**
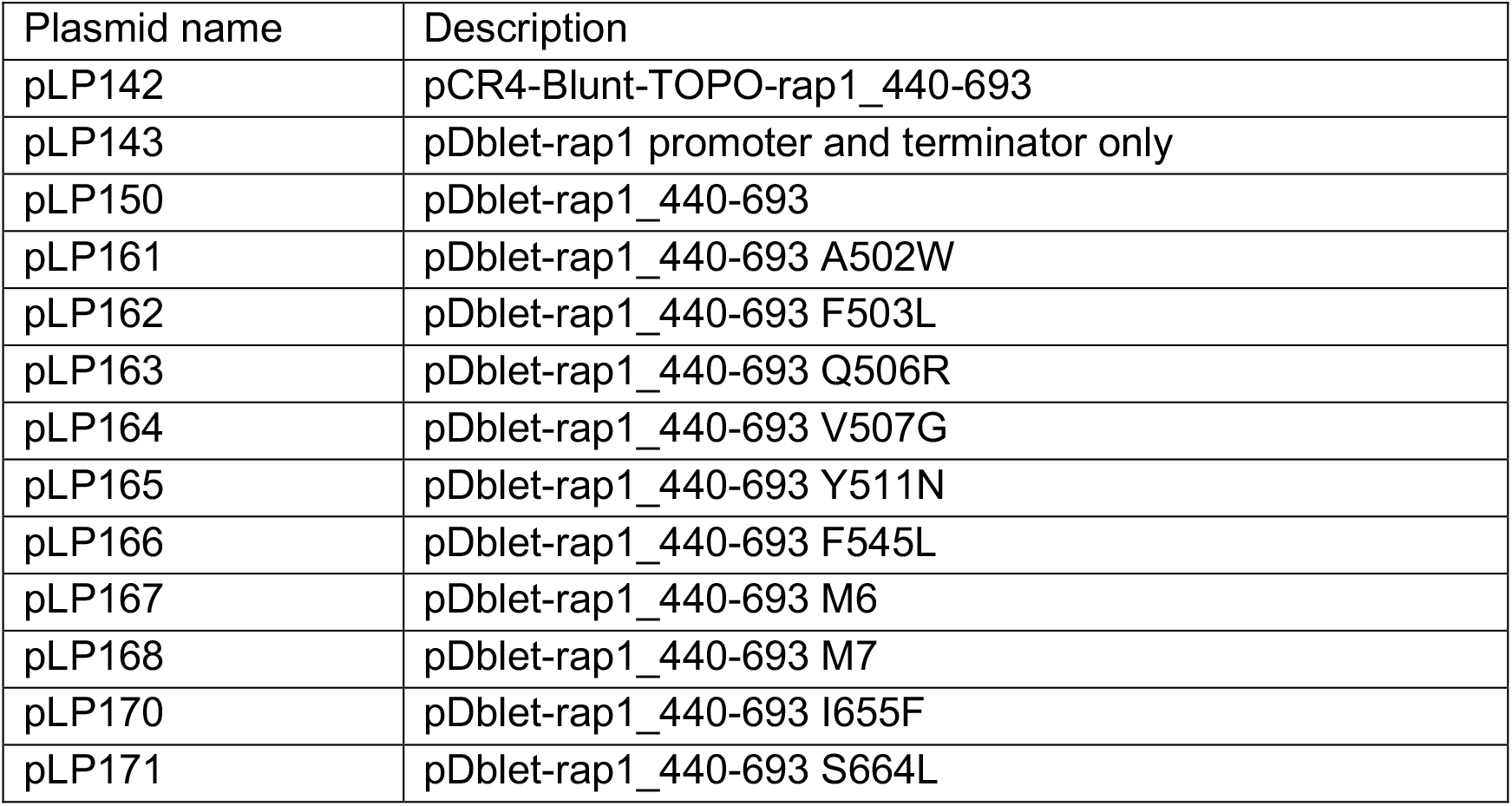
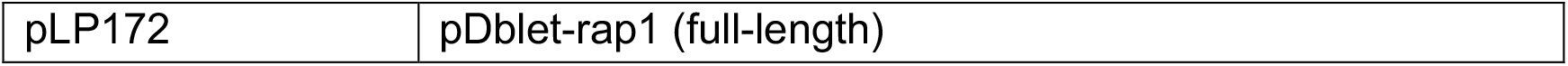
Plasmids used in this study.

**Table S4.**
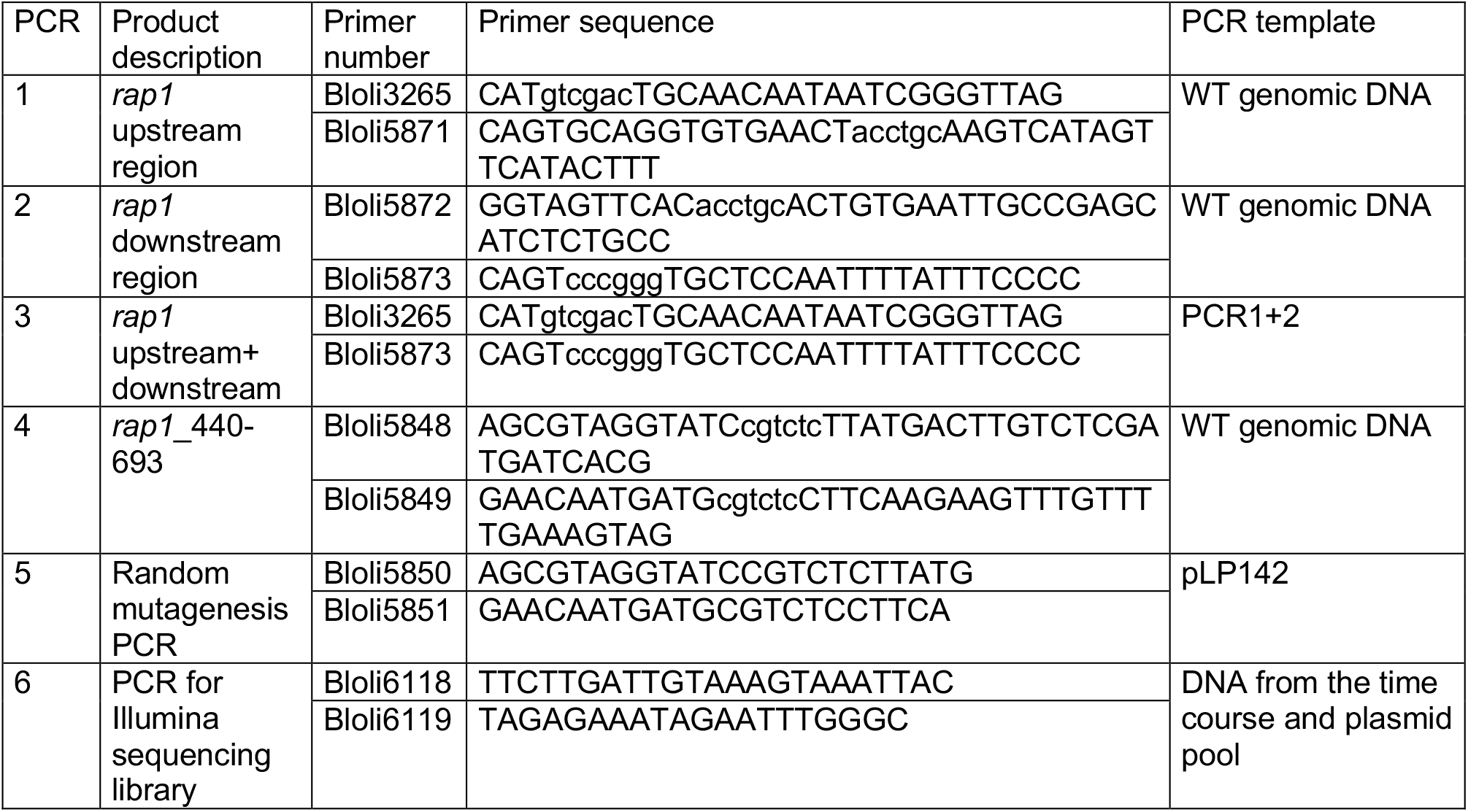
Oligos and templates used to construct mutagenesis library and generate PCR products for sequencing.

**Table S5.** Number of reads generated for each sequencing library. The Mut_lib_plasmid sample is the library purified from *E. coli* prior to introduction into fission yeast and ABC_R0 is the fission yeast R0 sample after recovery from plasmid transformation. A, B and C refer to the biological replicates and R1 to R5 to the rounds of arrest and return to growth.

| Sample name | Total reads |
| --- | --- |
| Mut_lib_plasmid | 714352 |
| ABC_R0 | 382374 |
| A_R1 | 751714 |
| A_R2 | 529933 |
| A_R3 | 813649 |
| A_R4 | 681322 |
| A_R5 | 620881 |
| B_R1 | 732036 |
| B_R2 | 560009 |
| B_R3 | 833834 |
| B_R4 | 763065 |
| B_R5 | 815709 |
| C_R1 | 873514 |
| C_R2 | 1074170 |
| C_R3 | 736946 |
| C_R4 | 777766 |
| C_R5 | 817761 |

## References

Armanios M (2009) Syndromes of telomere shortening. Annual review of genomics and human genetics 10: 45–61

Artandi SE, DePinho RA (2010) Telomeres and telomerase in cancer. Carcinogenesis 31: 9–18

Bae NS, Baumann P (2007) A RAP1/TRF2 complex inhibits nonhomologous end-joining at human telomeric DNA ends. Mol Cell 26: 323–334

Bahler J, Wu JQ, Longtine MS, Shah NG, McKenzie A 3rd,, Steever AB, Wach A, Philippsen P, Pringle JR (1998) Heterologous modules for efficient and versatile PCR-based gene targeting in Schizosaccharomyces pombe. Yeast 14: 943–951

Baumann P, Cech TR (2001) Pot1, the putative telomere end-binding protein in fission yeast and humans. Science 292: 1171–1175

Bilaud T, Brun C, Ancelin K, Koering CE, Laroche T, Gilson E (1997) Telomeric localization of TRF2, a novel human telobox protein. Nat Genet 17: 236–239

Bolger AM, Lohse M, Usadel B (2014) Trimmomatic: a flexible trimmer for Illumina sequence data. Bioinformatics 30(15): 2114–2120

Broccoli D, Smogorzewska A, Chong L, de Lange T (1997) Human telomeres contain two distinct Myb-related proteins, TRF1 and TRF2. Nat Genet 17: 231–235

Brun C, Dubey DD, Huberman JA (1995) pDblet, a stable autonomously replicating shuttle vector for Schizosaccharomyces pombe. Gene 164: 173–177

Chen Y, Rai R, Zhou ZR, Kanoh J, Ribeyre C, Yang Y, Zheng H, Damay P, Wang F, Tsujii H et al (2011) A conserved motif within RAP1 has diversified roles in telomere protection and regulation in different organisms. Nat Struct Mol Biol 18: 213–221

Chikashige Y, Hiraoka Y (2001) Telomere binding of the Rap1 protein is required for meiosis in fission yeast. Curr Biol 11: 1618–1623

Chikashige Y, Yamane M, Okamasa K, Tsutsumi C, Kojidani T, Sato M, Haraguchi T, Hiraoka Y (2009) Membrane proteins Bqt3 and -4 anchor telomeres to the nuclear envelope to ensure chromosomal bouquet formation. J Cell Biol 187: 413–427

Chong L, van Steensel B, Broccoli D, Erdjument-Bromage H, Hanish J, Tempst P, de Lange T (1995) A human telomeric protein. Science 270: 1663–1667

Cooper JP, Nimmo ER, Allshire RC, Cech TR (1997) Regulation of telomere length and function by a Myb-domain protein in fission yeast. Nature 385: 744–747

Crabbe L, Cesare AJ, Kasuboski JM, Fitzpatrick JA, Karlseder J (2012) Human telomeres are tethered to the nuclear envelope during postmitotic nuclear assembly. Cell Rep 2: 1521–1529

De Lange T (2005) Telomere-related genome instability in cancer. Cold Spring Harb Symp Quant Biol 70: 197–204

Ferreira MG, Cooper JP (2001) The fission yeast Taz1 protein protects chromosomes from Ku-dependent end-to-end fusions. Mol Cell 7: 55–63

Fujita I, Tanaka M, Kanoh J (2012) Identification of the functional domains of the telomere protein Rap1 in Schizosaccharomyces pombe. PLoS One 7: e49151

Hediger F, Neumann FR, Van Houwe G, Dubrana K, Gasser SM (2002) Live imaging of telomeres: yKu and Sir proteins define redundant telomere-anchoring pathways in yeast. Curr Biol 12: 2076–2089

Houghtaling BR, Cuttonaro L, Chang W, Smith S (2004) A dynamic molecular link between the telomere length regulator TRF1 and the chromosome end protector TRF2. Curr Biol 14: 1621–1631

Hu C, Inoue H, Sun W, Takeshita Y, Huang Y, Xu Y, Kanoh J, Chen Y (2019) Structural insights into chromosome attachment to the nuclear envelope by an inner nuclear membrane protein Bqt4 in fission yeast. Nucleic Acids Res 47: 1573–1584

Julin B, Shui I, Heaphy CM, Joshu CE, Meeker AK, Giovannucci E, De Vivo I, Platz EA (2015) Circulating leukocyte telomere length and risk of overall and aggressive prostate cancer. British journal of cancer 112: 769–776

Kanoh J, Ishikawa F (2001) spRap1 and spRif1, recruited to telomeres by Taz1, are essential for telomere function in fission yeast. Curr Biol 11: 1624–1630

Kim JK, Liu J, Hu X, Yu C, Roskamp K, Sankaran B, Huang L, Komives EA, Qiao F (2017) Structural Basis for Shelterin Bridge Assembly. Mol Cell 68: 698–714 e695

Kim SH, Kaminker P, Campisi J (1999) TIN2, a new regulator of telomere length in human cells. Nat Genet 23: 405–412

Lan Q, Cawthon R, Shen M, Weinstein SJ, Virtamo J, Lim U, Hosgood HD 3rd,, Albanes D, Rothman N (2009) A prospective study of telomere length measured by monochrome multiplex quantitative PCR and risk of non-Hodgkin lymphoma. Clin Cancer Res 15: 7429–7433

Li B, Lustig AJ (1996) A novel mechanism for telomere size control in Saccharomyces cerevisiae. Genes Dev 10: 1310–1326

Li B, Oestreich S, de Lange T (2000) Identification of human Rap1: implications for telomere evolution. Cell 101: 471–483

Li H, Durbin R (2009) Fast and accurate short read alignment with Burrows-Wheeler transform. Bioinformatics 25(14): 1754–1760

Li H, Handsaker B, Wysoker A, Fennell T, Ruan J, Homer N, Marth G, Abecasis G, Durbin R (2009) The Sequence Alignment/Map format and SAMtools. Bioinformatics 25(16): 2078–2079

Liu D, Safari A, O’Connor MS, Chan DW, Laegeler A, Qin J, Songyang Z (2004) PTOP interacts with POT1 and regulates its localization to telomeres. Nat Cell Biol 6: 673–680

Loayza D, De Lange T (2003) POT1 as a terminal transducer of TRF1 telomere length control. Nature 423: 1013–1018

Lototska L, Yue JX, Li J, Giraud-Panis MJ, Songyang Z, Royle NJ, Liti G, Ye J, Gilson E, Mendez-Bermudez A (2020) Human RAP1 specifically protects telomeres of senescent cells from DNA damage. EMBO reports 21: e49076

Lynch SM, Major JM, Cawthon R, Weinstein SJ, Virtamo J, Lan Q, Rothman N, Albanes D, Stolzenberg-Solomon RZ (2013) A prospective analysis of telomere length and pancreatic cancer in the alpha-tocopherol beta-carotene cancer (ATBC) prevention study. Int J Cancer 133: 2672–2680

Machiela MJ, Hofmann JN, Carreras-Torres R, Brown KM, Johansson M, Wang Z, Foll M, Li P, Rothman N, Savage SA et al (2017) Genetic Variants Related to Longer Telomere Length are Associated with Increased Risk of Renal Cell Carcinoma. European urology 72: 747–754

Machiela MJ, Hsiung CA, Shu XO, Seow WJ, Wang Z, Matsuo K, Hong YC, Seow A, Wu C, Hosgood HD 3rd, et al (2015) Genetic variants associated with longer telomere length are associated with increased lung cancer risk among never-smoking women in Asia: a report from the female lung cancer consortium in Asia. Int J Cancer 137: 311–319

Machiela MJ, Lan Q, Slager SL, Vermeulen RC, Teras LR, Camp NJ, Cerhan JR, Spinelli JJ, Wang SS, Nieters A et al (2016) Genetically predicted longer telomere length is associated with increased risk of B-cell lymphoma subtypes. Hum Mol Genet 25: 1663–1676

Maciejowski J, Li Y, Bosco N, Campbell PJ, de Lange T (2015) Chromothripsis and Kataegis Induced by Telomere Crisis. Cell 163: 1641–1654

Maestroni L, Reyes C, Vaurs M, Gachet Y, Tournier S, Geli V, Coulon S (2020) Nuclear envelope attachment of telomeres limits TERRA and telomeric rearrangements in quiescent fission yeast cells. Nucleic Acids Res 48: 3029–3041

Marcand S, Pardo B, Gratias A, Cahun S, Callebaut I (2008) Multiple pathways inhibit NHEJ at telomeres. Genes Dev 22: 1153–1158

McKinney W (2010) Data Structures for Statistical Computing in Python. In: Proceedings of the 9th Python in Science Conference, Millman S.v.d.W.a.J. (ed.) pp. 51–56

Miller KM, Ferreira MG, Cooper JP (2005) Taz1, Rap1 and Rif1 act both interdependently and independently to maintain telomeres. EMBO J 24: 3128–3135

Miller KM, Rog O, Cooper JP (2006) Semi-conservative DNA replication through telomeres requires Taz1. Nature 440: 824–828

Miyoshi T, Kanoh J, Saito M, Ishikawa F (2008) Fission yeast Pot1-Tpp1 protects telomeres and regulates telomere length. Science 320: 1341–1344

Moretti P, Freeman K, Coodly L, Shore D (1994) Evidence that a complex of SIR proteins interacts with the silencer and telomere-binding protein RAP1. Genes Dev 8: 2257–2269

Ojha J, Codd V, Nelson CP, Samani NJ, Smirnov IV, Madsen NR, Hansen HM, de Smith AJ, Bracci PM, Wiencke JK et al (2016) Genetic Variation Associated with Longer Telomere Length Increases Risk of Chronic Lymphocytic Leukemia. Cancer epidemiology, biomarkers & prevention : a publication of the American Association for Cancer Research, cosponsored by the American Society of Preventive Oncology 25: 1043–1049

Palm W, de Lange T (2008) How shelterin protects mammalian telomeres. Annu Rev Genet 42: 301–334

Pan L, Hildebrand K, Stutz C, Thoma N, Baumann P (2015) Minishelterins separate telomere length regulation and end protection in fission yeast. Genes Dev 29: 1164–1174

Pardo B, Marcand S (2005) Rap1 prevents telomere fusions by nonhomologous end joining. EMBO J 24: 3117–3127

Pellatt AJ, Wolff RK, Torres-Mejia G, John EM, Herrick JS, Lundgreen A, Baumgartner KB, Giuliano AR, Hines LM, Fejerman L et al (2013) Telomere length, telomere-related genes, and breast cancer risk: the breast cancer health disparities study. Genes Chromosomes Cancer 52: 595–609

Pfeiffer V, Lingner J (2013) Replication of telomeres and the regulation of telomerase. Cold Spring Harbor Perspectives in Biology 5: a010405

Pickett HA, Cesare AJ, Johnston RL, Neumann AA, Reddel RR (2009) Control of telomere length by a trimming mechanism that involves generation of t-circles. EMBO J 28: 799–809

Pinzaru AM, Kareh M, Lamm N, Lazzerini-Denchi E, Cesare AJ, Sfeir A (2020) Replication stress conferred by POT1 dysfunction promotes telomere relocalization to the nuclear pore. Genes Dev 34: 1619–1636

Rode L, Nordestgaard BG, Bojesen SE (2016) Long telomeres and cancer risk among 95 568 individuals from the general population. International journal of epidemiology 45: 1634–1643

Rubio MA, Davalos AR, Campisi J (2004) Telomere length mediates the effects of telomerase on the cellular response to genotoxic stress. Exp Cell Res 298: 17–27

Sarthy J, Bae NS, Scrafford J, Baumann P (2009) Human RAP1 inhibits non-homologous end joining at telomeres. EMBO J 28: 3390–3399

Schindelin J, Arganda-Carreras I, Frise E, Kaynig V, Longair M, Pietzsch T, Preibisch S, Rueden C, Saalfeld S, Schmid B et al (2012) Fiji: an open-source platform for biological-image analysis. Nat Methods 9: 676–682

Schober H, Ferreira H, Kalck V, Gehlen LR, Gasser SM (2009) Yeast telomerase and the SUN domain protein Mps3 anchor telomeres and repress subtelomeric recombination. Genes Dev 23: 928–938

Sfeir A, Kabir S, van Overbeek M, Celli GB, de Lange T (2010) Loss of Rap1 induces telomere recombination in the absence of NHEJ or a DNA damage signal. Science 327: 1657–1661

Sfeir A, Kosiyatrakul ST, Hockemeyer D, MacRae SL, Karlseder J, Schildkraut CL, de Lange T (2009) Mammalian telomeres resemble fragile sites and require TRF1 for efficient replication. Cell 138: 90–103

Sievers F, Wilm A, Dineen D, Gibson TJ, Karplus K, Li W, Lopez R, McWilliam H, Remmert M, Soding J et al (2011) Fast, scalable generation of high-quality protein multiple sequence alignments using Clustal Omega. Mol Syst Biol 7: 539

Therizols P, Fairhead C, Cabal GG, Genovesio A, Olivo-Marin JC, Dujon B, Fabre E (2006) Telomere tethering at the nuclear periphery is essential for efficient DNA double strand break repair in subtelomeric region. J Cell Biol 172: 189–199

Wang RC, Smogorzewska A, de Lange T (2004) Homologous recombination generates T-loop-sized deletions at human telomeres. Cell 119: 355–368

Yang CW, Hsieh MH, Sun HJ, Teng SC (2021) Nuclear envelope tethering inhibits the formation of ALT-associated PML bodies in ALT cells. Aging (Albany NY) 13: 10490–10516

Ye JZ, Hockemeyer D, Krutchinsky AN, Loayza D, Hooper SM, Chait BT, de Lange T (2004) POT1-interacting protein PIP1: a telomere length regulator that recruits POT1 to the TIN2/TRF1 complex. Genes Dev 18: 1649–1654

Zhong Z, Shiue L, Kaplan S, de Lange T (1992) A mammalian factor that binds telomeric TTAGGG repeats in vitro. Mol Cell Biol 12: 4834–4843

